# Epigenetic transcriptional reprogramming by WT1 mediates a repair response during podocyte injury

**DOI:** 10.1101/2020.02.18.954347

**Authors:** Sandrine Ettou, Youngsook L. Jung, Tomoya Miyoshi, Dhawal Jain, Ken Hiratsuka, Valerie Schumacher, Mary E. Taglienti, Ryuji Morizane, Peter J. Park, Jordan A. Kreidberg

## Abstract

In the context of human disease, the mechanisms whereby transcription factors reprogram gene expression in response to injury are not well understood. This is particularly true in kidney podocytes, injury to which is the common and initial event in many processes that lead End Stage Kidney Disease. WT1 is a master regulator of gene expression in podocytes, binding nearly all genes known to be crucial for maintenance of the glomerular filtration barrier. Here, using purified populations of podocytes and glomeruli, we investigated WT1-mediated transcriptional reprogramming during the course of podocyte injury. Using the Adriamycin murine model of Focal Segmental Glomerulosclerosis, we discovered that podocyte injury led to increased intensity of WT1 binding and to the acquisition of new WT1 binding sites, both at previously identified target genes and at newly bound target genes, providing mechanistic insight on the transcriptional response to injury. We also observed a previously unrecognized transient increase in expression of WT1 target genes in both mice and human kidney organoids. Together, these features appear to constitute an attempt to repair the glomerular filtration barrier after podocyte injury. At later stages of injury, when proteinuria became severe, there was greatly decreased WT1 binding to most target genes. Furthermore, WT1 appeared to be required to maintain active chromatin marks at its target genes. These active marks were converted to repressive marks after loss of WT1 or Adriamycin-induced injury. This response to injury suggests that there may be a potential window of opportunity for repairing podocyte injury as a treatment for glomerular disease in humans.

## INTRODUCTION

Focal Segmental Glomerulosclerosis (FSGS) is among the most debilitating and least treatable forms of chronic kidney disease and often leads to End Stage Kidney Disease, requiring dialysis and/or transplantation. Podocytes are highly differentiated cells that maintain the glomerular filtration barrier (GFB) through the extension of foot proceses that interdigitate with foot processes of adjacent podocytes, thereby assembling a scaffold that supports a network of capillaries within each glomerulus. In most types of FSGS, podocyte injury is the incipient event^1^, characterized by foot process effacement and podocyte detachment, resulting in loss of the GFB and severe proteinuria^2^. Several proteins are implicated in maintaining podocyte structural organization, including Synaptopodin, Nephrin and Podocin. One common characteristic of glomerular injury is the decreased abundance of key proteins that maintain the GFB, suggesting that transcriptional regulation of genes encoding these proteins has an important role in the pathogenesis of glomerular disease.

Our previous study and others identified WT1 as one of the most upstream transcription factors (TF) regulating gene expression in podocytes^3, 4^ and one of the earliest known markers of podocytes during kidney development^5^. Decreased expression of *WT1* and mutations in *WT1* gene have been described in several forms of glomerular disease^6–10^. Most human nephrotic syndrome genes have been identified as WT1 targets, including *NPHS1, NPHS2*, and *INF2* ^4^. However, the mechanism whereby WT1 regulates gene expression during the initiation and progression of glomerular disease remains unknown.

In the present study, we focused on deciphering the transcriptional mechanisms through which WT1 regulates podocyte gene expression during injury. Genome-wide analysis of both WT1 DNA occupancy and podocyte gene expression during the course of Adriamycin-mediated injury revealed a transient increase in the number and binding intensity (defined as peak height in ChIP-seq data sets) of WT1-bound sites as well as an increase in the expression of crucial podocyte genes at early stages of injury, that may reflect an attempt to repair podocytes. We demonstrated that WT1 is required to maintain active chromatin marks at podocyte genes, and that podocyte injury leads to the conversion from active to repressive histone modifications at *Nphs2* and *Synpo*. Taken together, this study provides strong evidence that, during injury, podocyte gene expression is subject to transcriptional reprogramming under the direct control of WT1, indicating that podocytes possess an intrinsic repair program, acting at the level of gene expression.

## RESULTS

### Epigenetic regulation of podocyte gene expression by WT1

WT1 has been identified as a key regulator of podocyte gene expression^3, 4^ and WT1 target genes are crucial for maintaining glomerular filtration barrier^4, 11^. Two WT1 target genes were studied to elucidate the transcriptional response to injury. *Nphs2* encodes Podocin, an essential component of the slit diaphragm and *Synpo* encodes Synaptopodin, an actin-associated protein important for maintaining the cytoskeleton integrity. To directly demonstrate WT1-dependent gene expression, *Wt1* was conditionally inactivated in podocytes^12^ of adult *Nphs2-CreERT2, WT1fl/fl* mice, leading to massive proteinuria (Fig. 1a). Kidneys appeared pale (Fig. 1b) with H&E and PAS staining showing protein casts, mesangial expansion and dilated tubules (Fig. 1c). WT1, Podocin and Synaptopodin transcript and protein levels were greatly reduced (Fig. 1d, e).

**Fig. 1.**
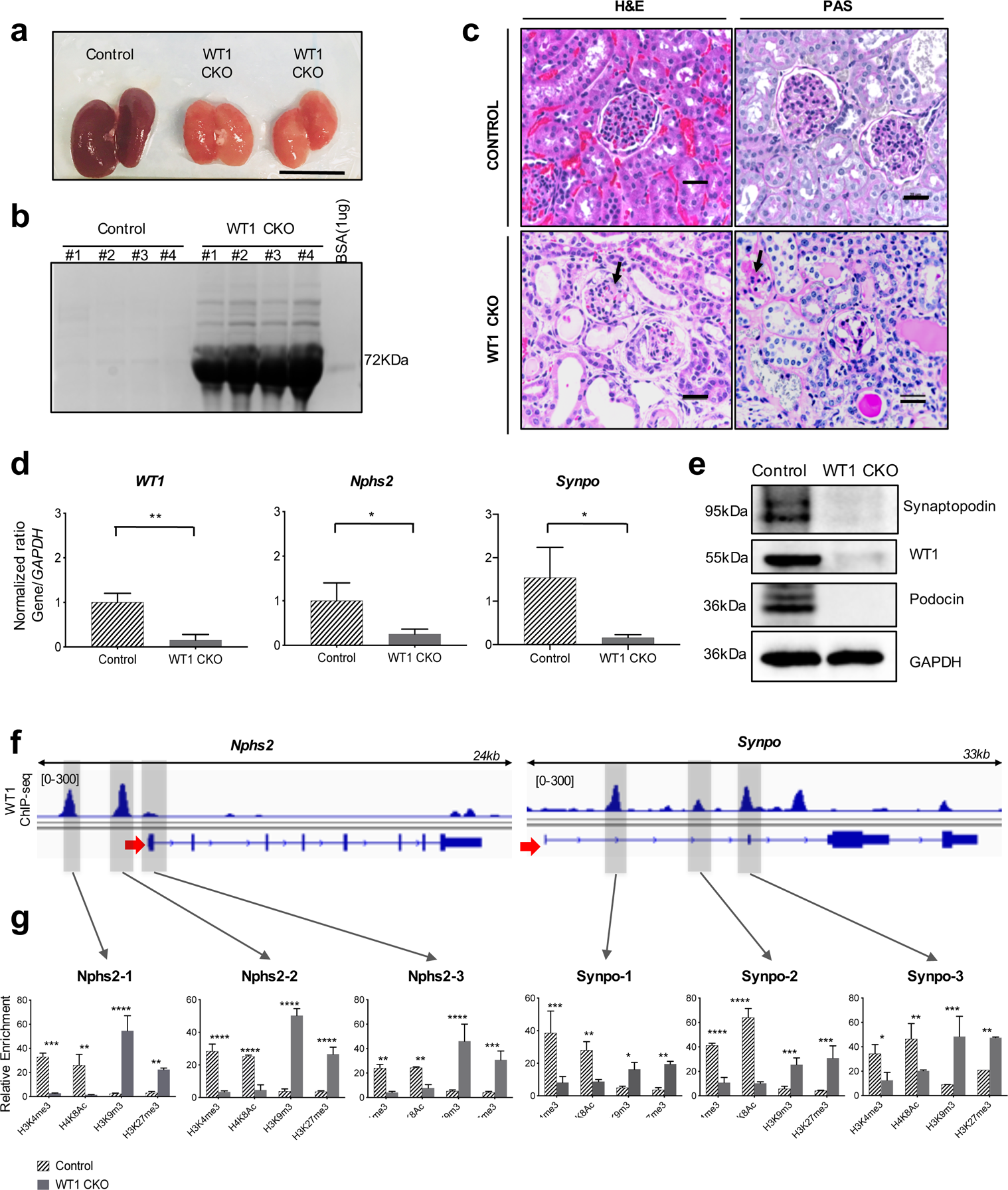
WT1 controls chromatin remodeling at *Nphs2* and *Synpo* genes in mice. (**a**) *WT1^flox/flox^/iNphs2-cre* mice exhibit smaller and pale kidneys compared to control (n=3) at D14 after tamoxifen injection. Scale bar 1cm. (**b**) Coomassie stains gel of 5 μl urine from *WT1^flox/flox^* (control) mice and *WT1^flox/flox^/iNphs2-cre* (WT1 CKO) mice (control: BSA) Each lane represents a single mouse. (**c**) Representative histological images of control and WT1 CKO kidneys by hematoxylin and eosin (H&E) and periodic acid-Schiff (PAS) at D14 after tamoxifen injections. Original magnification, x60, scale bars: 20μm. Black arrows: mesengial expansion. (**d**) RT-qPCR of *Wt1, Nphs2* and *Synpo* from control and WT1 CKO mice. Bars represent means and error bars ± SEMs. *\*\*P*<0.01; *\*P*<0.05 (n=3). (**e**) Representative western blot (of three independent experiments) reflecting WT1 expression from control and WT1 CKO mice at D14 after tamoxifen injections. (**f**) IGV plots of *Nphs2* and *Synpo* genes for WT1 ChIP-seq showing WT1 binding sites (gray highlighted boxes) in uninjured podocytes: Nphs2-1, Nphs2-2, Nphs2-3, Synpo-1, Synpo-2 and Synpo-3. (**g**) histone direct ChIP-qPCR from FACS-isolated podocytes from control and WT1 CKO mice 14 days after tamoxifen injections, using active histones marks (H3K4m3, H4K8ac) and repressive histones marks (H3K9me3 and H3K27me3). **** *P<0.0001*, *** *P<0.001*, ** *P<0.01*, *\* P<0.05* (Multiple *t*-test with FDR determined using the two-stage linear step-up procedure of Benjamini, Krieger and Yekutieli) compared to control mice.

Tissue specific TFs activate gene expression in part by promoting histone modifications that maintain open chromatin, such as H3K4me3 and H4K8ac. We used FACS-isolated podocytes to analyze the effect of WT1 inactivation on histone modifications during the course of injury at previously defined WT1 binding sites at the *Nphs2* and *Synpo* genes^4^, here identified as Nphs2-1, Nphs2-2, Nphs2-3, Synpo-1, Synpo-2 and Synpo-3. Nphs2-1 and Nphs2-2 are located upstream of the promoter and are putative enhancers. Nphs2-3 is at the transcriptional start site (TSS). Synpo-1 and Synpo-2 are located in intronic regions and Synpo-3 overlaps the second exon (Fig. 1f). H3K4me3 and H4K8ac were greatly reduced after inactivation of WT1 at these sites (Fig. 1g) and the repressive histones marks H3K9me3 and H3K27me3 were increased (Fig. 1g), confirming that histones modification complexes that maintain the active chromatin state also inhibit the placement of repressive marks on histones^13^. Similar results were observed *in vitro* with immortalized podocytes (Supplementary Fig. 1a, b), demonstrating that WT1 has a crucial role in maintaining the open chromatin state at its target genes in podocyte.

### Transient increase of podocyte gene expression in ADR-injured mice and human kidney organoids

To analyze WT1-mediated transcriptional reprogramming during the course of injury, we used the Adriamycin (ADR) model for podocyte injury, a well-recognized murine model for FSGS^14, 15^. Two different strains of mice were used in this study: *mTmG-Nphs2cre* mice that are less sensitive to ADR, from which podocytes are isolated by FACS, and BALB/cJ, a prototypical highly ADR-sensitive strain^16^. To determine the time course of ADR-induced podocyte injury, we first analyzed the level of proteinuria of *mTmG-Nphs2cre* and BALB/cJ mice treated with either ADR or phosphate buffered saline (PBS) as a control vehicle. *mTmG-Nphs2cre* mice required a higher dose and a second injection to develop maximal proteinuria after two weeks (Fig. 2a), whereas ADR induced proteinuria in BALB/cJ mice over a one-week period (Supplementary Fig. 2a). The expression of WT1 has previously been shown to decrease after the onset of glomerular disease^17, 18^. However, in BALB/cJ isolated glomeruli, we observed a several-fold increase in *Wt1* expression after ADR treatment, before *Wt1* fell to low levels (Supplementary Fig. 2b). Concomitant with the transient increase in *Wt1*, *Nphs2* and *Synpo* also dramatically increased before falling to nearly undetectable levels (Supplementary Fig. 2b). Although, in *mTmG-Nphs2cre* FACS-isolated podocytes, *Wt1* and *Nphs2* levels did not increase to the extent observed in BALB/cJ after ADR, *Synpo* did show an over two-fold increase, suggesting that while less dramatic, there also appeared to have been transcriptional reprogramming in these mice (Fig. 2b). In *mTmG-Nphs2cre* mice, WT1 protein levels fell at day 9 after ADR (D9). Podocin showed a slight increase whereas Synaptopodin showed an over three-fold increase at Day 5 after Adriamycin (D5), then falling dramatically (Fig. 2c). By immunofluorescent detection, WT1 was also present in podocyte nuclei until D5 in *mTmG-Nphs2cre* and D3 in BALB/cJ, after which it was decreased (Fig. 1d and Supplementary Fig. 2c).

**Fig. 2.**
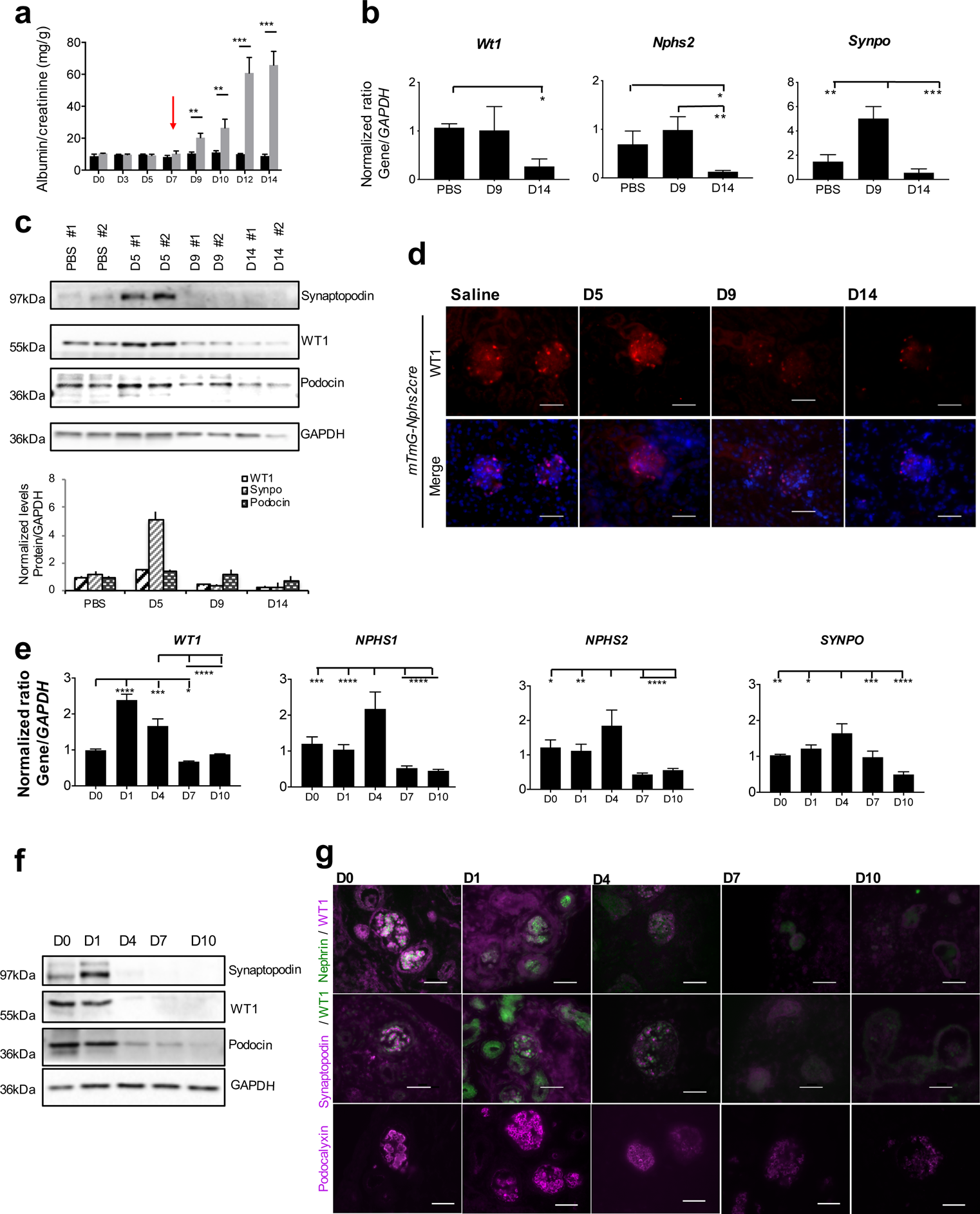
Transient increase in the expression of key podocyte genes in ADR-injured mice and human organoids. (a) Quantification of albumin/creatinine level during the course of ADR injury from *mTmG-Nphs2cre* mice injected twice with 18mg/kg of ADR (grey bars) or PBS (black bars) at one-week intervals (second injection indicated by red arrow). Bars represent means and error bars ± SEMs. *\*\*\*P*<0.001*; **P*<0.01 (n=3 replicates). (b) RT-qPCR of *Wt1*, *Nphs2* and *Synpo* from *mTmG-Nphs2cre* FACS-isolated podocytes during injury. One-way ANOVA with Tukey’s multiple comparisons test were used. \*\*\**P*<0.001; \*\**P*<0.01; *\*P*<0.05 (n=3 replicates). (**c**) Representative western blot (of three independent experiments performed) reflecting podocyte protein levels during the course of ADR-induced injury from *mTmG-Nphs2cre* isolated glomeruli. Each lane represents a single mouse. Lower panel: quantification of western blot based on n=3. (**d**) Immunofluorescent staining of WT1 (red) in *mTmG-Nphs2cre* mice in glomeruli. Scale bar 50µM. (**e**) RT-qPCR of *WT1*, *NPHS1, NPHS2* and *SYNPO* from human kidney organoids treated with 10μM of ADR during 1, 4, 7 and 10 days. ANOVA with Tukey’s multiple comparisons test were used. **** *P*<0.0001, *** *P*<0.001, ** *P*<0.01, *\* P*<0.05 (n=3). (**f**) Representative western blot (of three independent experiments performed) reflecting podocyte protein levels during the course of ADR-induced injury from human kidney organoids. (**g**) Immunofluorescent localization of WT1, NPHS1 and Podocalyxin at D0, D1, D4, D7 and D10 post treatment with ADR. Scale bars: 50μm.

We next investigated the effect of ADR on organoids derived from human embryonic stem cells. In addition to *WT1, NPHS2 and SYNPO,* we also examined *NPHS1,* encoding the slit diaphragm protein Nephrin, because of its importance in genetic kidney disease. We observed similar transient increases in WT1 and target genes (Figure 2e,f,g). Even though transcript levels continued to be expressed through D4 (Fig. 2e), protein levels for WT1, Synaptopodin, Nephrin (immunofluorescence only) and Podocin were greatly decreased starting from D4 by either western blot or immunofluorescent detection (Fig. 2f,g). Podocalyxin, a glycocalyx sialoprotein located at the apical and lateral surface of podocytes could be detected, albeit at lower levels, through D10, confirming their presence of podocytes throughout the injury process (Fig. 2g). Therefore, human podocytes also respond to injury by transiently increasing expression of WT1 and target genes, thus validating human kidney organoids as a model to study glomerular injury. Since protein levels did not precisely overlap with maximal RNA levels, there may also be translational regulation affecting their expression during injury. Nevertheless, the decrease in WT1 and target gene expression demonstrated significant transcriptional reprogramming during the course of the injury.

### Dynamics of WT1 occupancy during ADR-induced injury

The overall level of WT1 in podocytes may not be representative of its binding at specific enhancers or TSSs. The pattern of WT1 binding was indeed distinct at each site (shown in figure 3d) for *Nphs2* and *Synpo* genes during injury (Fig. 3a). In *mTmG-Nphs2cre* mice, the most significant changes were increased binding at Nphs2-1 at D10 and decreased binding at Nphs2-3 after D5. Binding at all three *Synpo* sites transiently increased, before falling to levels below that observed in uninjured mice, correlating with gene expression. This response is more dramatic in BALB/cJ mice, consistent with their greater sensitivity to ADR (Supplementary Fig. 2d). An important point emerges from this study that may be generalizable to transcription factor biology in general and particularly in response to disease. WT1 binding at distinct enhancers and promoters does not simply reflect the overall level of WT1, but may differ between individual binding sites. Recent studies have demonstrated redundancy among multiple enhancers affecting expression of individual genes during development^19^. In contrast, our results suggest that enhancers may make distinct contributions to gene expression during the response to injury.

**Fig. 3.**
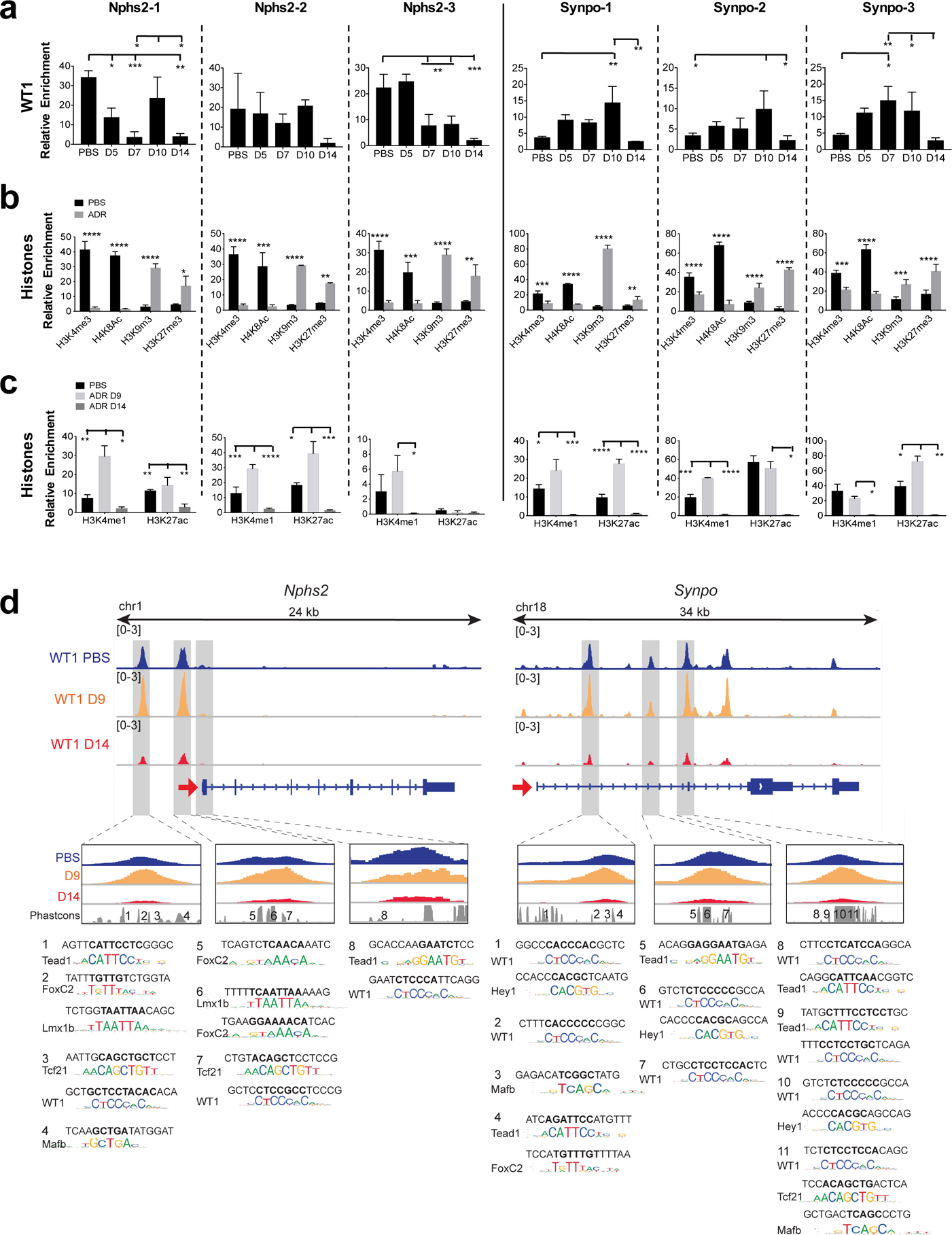
Effect of WT1 dynamic binding on chromatin remodeling during ADR-induced injury in *mTmG-Nphs2cre* mice. (**a**) WT1 dynamic binding at three binding sites on *Nphs2* and *Synpo* genes measured by WT1 direct ChIP-qPCR from isolated glomeruli from *mTmG-Nphs2cre* mice (n=3). ANOVA with Tukey’s multiple comparisons test were used. \*\*\**P*<0.001; \*\**P*<0.01; *\*P<*0.05. (**b**) Histone direct ChIP-qPCR at D14 from FACS-isolated podocytes from *mTmG-Nphs2cre* mice, using active histones marks (H3K4me3, H4K8ac) and repressive histones marks (H3K9me3 and H3K27me3) (n=3). **** *P<*0.0001, *** *P<*0.001, ** *P<*0.01, *\* P<*0.05 (Multiple *t*-test with FDR determined using the two-stage linear step-up procedure of Benjamini, Krieger and Yekutieli) compared to uninjured mice. (**c**) Histone direct ChIP-qPCR at D9 and D14 from FACS-isolated podocytes from *mTmG-Nphs2cre* mice using active enhancer marks (H3K4me1, H3K27ac) (n=3). ANOVA with Tukey’s multiple comparisons test were used. **** *P<*0.0001, \*\*\**P*<0.001; \*\**P*<0.01; *\*P<*0.05. (**d**) WT1 ChIP-seq profiles at *Nphs2* and *Synpo* genes during injury and predicted TF binding sites. Upper panels: IGV plots of WT1 binding at different sites during injury (uninjured: blue, D9: orange, D14: red). Red arrows show TSSs and direction of transcription. Gray highlighted boxes indicate WT1 binding sites shown in Fig. 1f. Lower panels: TF motifs identified within the numbered conserved elements.

### Chromatin remodeling during ADR-induced injury

As WT1 occupancy maintains open chromatin and ADR results in loss of WT1 binding at target genes (Fig. 1f), we interrogated histone modifications at WT1 binding sites after ADR treatment (Fig. 3b). All WT1 binding sites were converted to closed chromatin state at D14 after ADR injury (Fig. 3b). H3K4me1, that marks active enhancers and promoters, was increased at WT1 binding sites at D9 (Fig. 3c), correlating with increased binding of WT1. H3K27ac, that marks active enhancers, was also increased at D9, except at the Nphs2-3 site present within a TSS (Fig. 3c). Similar results were observed *in vitro* (Supplementary Fig. 1c,d). Thus, ADR injury results in a conversion from an open to closed chromatin state at WT1 target genes.

### Genome wide dynamics of WT1 binding during injury

To identify general mechanisms through which WT1 regulates gene expression during the course of ADR-induced podocyte injury, we performed WT1 ChIP-seq using isolated glomeruli from *mTmG-Nphs2cre* and BALB/cJ mice and RNA-seq using FACS-isolated podocytes from *mTmG-Nphs2cre* mice obtained at the onset of proteinuria (D9) and at a point of maximal proteinuria (D14), (Fig.1a). In *mTmG-Nphs2cre* mice, the global number of WT1 bound sites increased from 23,163 in control mice to 31,639 at D9 before falling dramatically to 6,567 binding sites at D14 (Fig. 4a). 11,266 binding sites were uniquely present at D9 (Fig. 4a). Many of these sites were present at genes already bound in uninjured podocytes (n=2,839; Fig. 4a and examples in Supplementary Fig. 3a). However, some of these sites were found at 1,245 genes not bound in control or D14 (Fig. 4b and examples in Supplementary Fig. 3b). Thus at D9, WT1 both bound additional sites at previously identfiied target genes, and acquired new target genes (Fig. 4b). Similar to D14 *mTmG-Nphs2cre* mice, proteinuric BALB/cJ mice also lost many WT1 binding sites at D7 after ADR (Supplementary Fig. 4a).

**Fig. 4.**
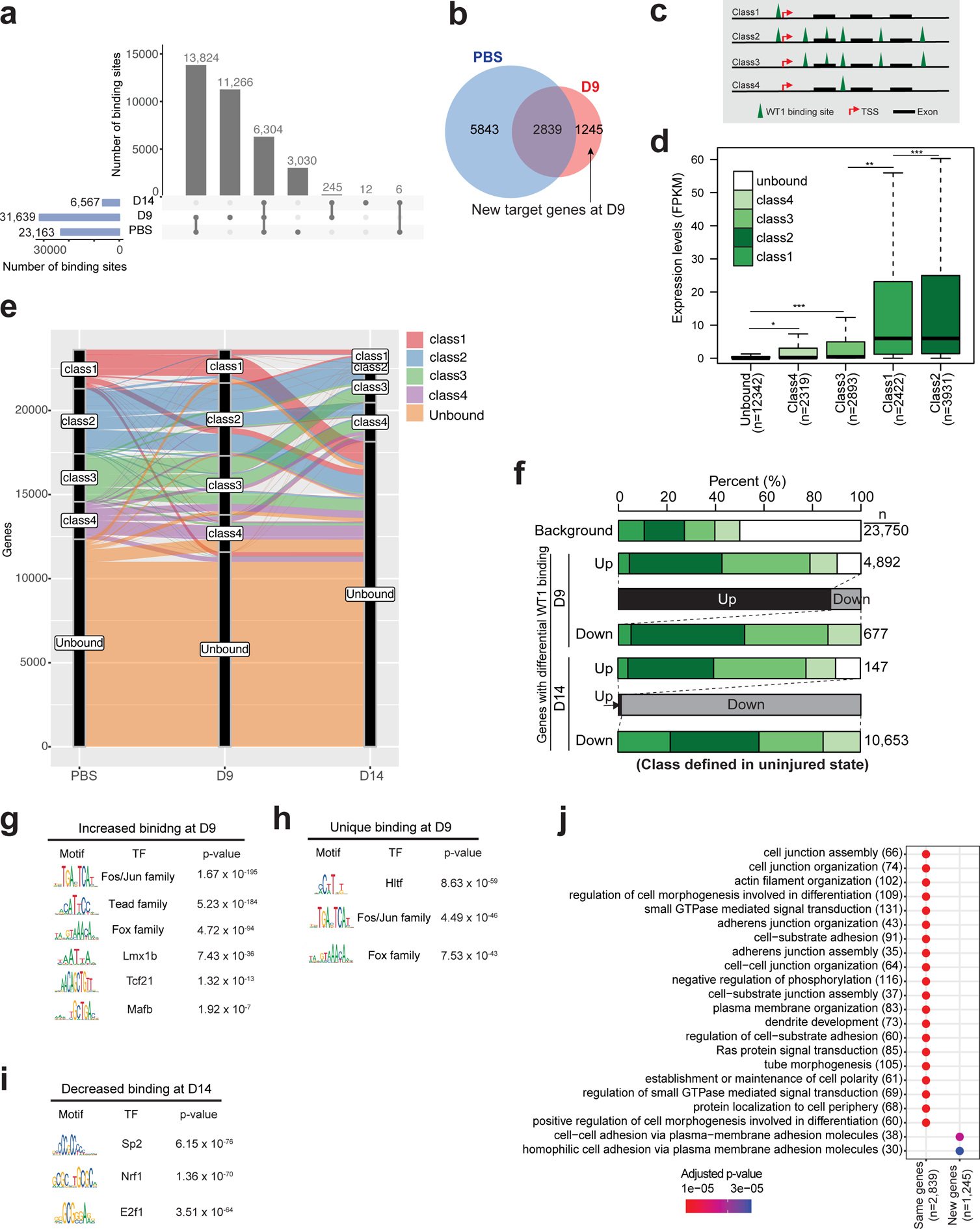
Dynamics of WT1 binding during the injury response in *mTmG-Nphs2cre* mice. (**a**) Number of WT1 binding sites during ADR injury. Grey bar plots represent the number of binding sites common to each condition. Number of total binding sites increased at Day 9 (D9) after ADR before decreasing at D14 (blue bar plot). (**b**) Venn diagram: The 11,266 WT1 binding sites (Figure 2A) unique to D9 are distributed between 1245 genes uniquely bound at D9, and 2839 genes that were also bound in control but at distinct sites (pink circle). 5384 genes were only bound in control. (**c**) Representation of the WT1 target gene classes. Class 1: genes having a single WT1 peak within the promoter region (defined as +/-1kb of the TSS); class 2: having multiple binding sites within a 500 kb region of the TSS including one at the promoter; class 3: having multiple binding sites but no binding at the promoter, class 4: having a single binding site that is not at the promoter. Unbound: having no binding within 500kb of the TSS. (**d**) Expression levels for each WT1 target gene class in FACS-isolated podocytes from control *mTmG-Nphs2cre* mice. *P*-values from Student’s t-test. *** *P*<0.001, ** *P*<0.01, *\* P*<0.05. In all expression bar graphs, the box shows the range of expression of the middle two quartiles, with the black line indicating median expression level. (**e**) Alluvial diagram showing gene class changes during the course of injury: class 1 (pink), class 2 (blue), class 3 (green), class 4 (purple) and unbound class (orange). Y-axis represents the number of genes per class, and X-axis the injury time points. (**f**) Proportion of each gene class for the genes associated with significant changes in WT1 binding intensity after ADR injury. Gene classes are based on the WT1 binding status in uninjured podocytes. Background indicates the distribution of gene classes for all genes in uninjured condition. The number in parenthesis for the last four columns represents the number of genes with WT1 binding sites that significantly changed during the course of injury. Significant changes in WT1 binding after ADR were mainly observed in classes 2 and 3, defined as containing multiple WT1 binding sites in uninjured podocytes (*P<0.001*, Fisher’s exact test compared to the expectation based on the background distributions of gene classes in uninjured podocytes). (**g)** TF Motifs frequently found near sites where WT1 binding increased at D9 compared to uninjured podocytes. (**h**) TF Motifs frequently found near WT1 binding sites uniquely present at D9. (**i**) TF Motifs frequently found near sites where WT1 binding decreased at D14 compared to uninjured podocytes. (**j**) GO terms based on unique WT1 binding sites at D9. New genes: GO terms based on 1,245 genes as shown in Fig. 4b; Same genes: GO terms based on 2,839 genes as shown in Fig. 4b.

Genome-wide, promoter regions were over-represented among WT1 binding sites (Supplementary figure 3a). The global distribution of WT1 binding sites did not change during injury in either strain of mice (Supplementary Fig. 4b and 5a). However, the intensity of WT1 binding changed over the course of injury. In *mTmG-Nphs2cre* mice, 93% of differentially bound sites increased at D9 whereas almost all binding sites significantly decreased at D14 (Supplementary Fig. 5b). In BALB/cJ mice, WT1 binding significantly changed (FDR<0.05) for 52% of the sites during injury, 85% of the differential WT1 binding sites decreased in intensity (Supplementary Fig. 4c). We observed differences in the functional distribution among these sites. Most notably, WT1 bound sites that increased in intensity after injury were primarily found in introns (50%) in *mTmG-Nphs2cre* mice, suggesting that WT1 bound additional intronic enhancer sites during the injury response (Supplementary Fig. 5b). These results demonstrate a process whereby, in the early stages of injury, WT1 acquires new binding sites and increases the intensity of its binding at previously bound sites.

### Dynamics of WT1 target gene classes during the injury response

We previously defined two classes of WT1 target genes, based on WT1 binding patterns (8): class 1 genes having a single WT1 binding site at the transcriptional start site (TSS +/-1kbp) and class 2 genes having multiple binding sites within a 500 kb region of the TSS including at the TSS. To these, we add class 3 genes having multiple WT1 binding sites but not at the TSS, and class 4 genes that have a single binding site not within 1kb of a TSS (Fig. 4c). Unbound genes are defined as not having a WT1 binding site within 500 kb of the TSS. Comparable to our previous findings, class 1/2 genes had a higher range of expression levels in uninjured podocytes. Classes 3/4, while expressed at lower levels than class 1/2 genes, were significantly more highly expressed than unbound genes (Fig. 4d), Thus, WT1 binding is a major determinant of gene expression in podocytes, and binding at the TSS is particularly important.

A large number of genes changed their class designation during the course of injury. Many genes not bound in uninjured podocytes became transiently bound at D9. A small number of genes changed from class 3/4 to class 1/2 at D9, and returned to class 3/4 or unbound at D14 (Fig. 4e). Class 2 and 3 genes showed the greatest changes in WT1 binding, the majority increasing at D9 and decreasing at D14 (*P*<0.001; Fig. 4f, and Supplementary Fig. 4e). Furthermore, in both strains of mice, many class 1/2 genes became unbound at D14 (Fig. 4e, Supplementary Fig. 4d). At D14, the number of genes with decreased or loss of WT1 binding (defined as either decreased number of binding sites, or binding intensity), greatly outnumbered bound genes (Fig. 4e,f). These analyses emphasize the importance of WT1 binding for the response to injury in podocytes.

### WT1 regulated transcriptional network

Eukaryote TFs generally act combinatorially to determine tissue-specific patterns of gene expression. Therefore, we examined TF motif enrichment near those WT1 binding sites whose intensity significantly increased at D9. This analysis highlighted motifs predicting FOX, LMX1B, TCF21 and MAFB as TFs potentially co-binding with WT1 (Fig. 4g), all well known to be important in podocytes^20–23^, suggesting that the response to injury involves the basic transcriptional machinery already present in podocytes. TEAD sites were also predicted, as were FOS/JUN sites, the former predicting a role for YAP/TAZ in regulating the injury response and the later suggesting a role for the AP-1 TF, described to confer protection from glomerulonephritis^24^. Near WT1 sites uniquely bound at D9, motifs predicting FOX TF co-binding were present, but not other podocyte TFs (Fig. 4h), suggesting that during the injury response, WT1 and FOX TFs may additionally activate a set of enhancers. Interestingly, motif analysis at sites where WT1 binding was decreased at D14 predicted an entirely different set of TFs, including SP2, NRF1 and E2F1/4 (Fig. 4i). NRF1 and E2F1/4 have been described as repressive TFs^25–27^, suggesting that by D14, a portion of WT1’s overall activity may be as part of a repressive complex involved in decreasing the expression of many target genes.

Predicted TF motifs at the *Nphs2* and *Synpo* genes are shown in Fig. 3d, (adapted from^4^). Interestingly, this analysis also indicates that individual enhancers may be bound by distinct groupings of TFs, suggesting that levels of gene expression reflect the integration of multiple enhancers, each of which might contribute differently to gene expression.

### Functional implications of WT1 dynamic binding

To understand the functional implications of WT1 dynamic binding, we performed RNA-seq analysis on control, D9 and D14 FACS-isolated podocytes from *mTmG-Nphs2-Cre* mice. GO analysis based on RNA-seq (Fig. 5a) and ChIP-seq (Supplementary Fig. 6a) data sets were largely consistent with each other in uninjured and D9 podocytes, emphasizing cytoskeletal organization and cell adhesion. Glomerular development was also identified as a GO term at D9, indicating that repair processes involved several genes implicated in formation of the glomerulus (Supplementary Fig. 6a). Interestingly, gene expression of, and binding of WT1 to, cytoskeletal and adhesion genes sets was decreased at D14, suggesting that by this point, repair processes in podocytes are greatly diminished (Fig. 5a and Supplementary Fig. 6b). GO analysis of the 1,245 genes uniquely bound at D9 (Fig. 4j) were similar to those identified as having increased intensity of WT1 binding at D9 (Supplementary Fig. 6b), indicating that the early response to injury largely involved amplification of the same pathways already operational in uninjured podocytes. Genes represented by GO terms related to RNA stability, nucleotide metabolism and splicing process showed decreased WT1 binding at D14 (Supplementary Fig. 6b), though their expression levels increased (Fig. 5a), further suggesting a potential repressive function for WT1 (examples of genes in Supplementary Fig. 6c,d). In BALB/cJ mice, GO analysis of genes at which binding increased after injury included regulation of protein processes, while genes at which WT1 binding decreased after injury included paraxial mesodermal development, a set that includes *Foxc2* and *Tead1* (Supplementary Fig. 4f,g).

**Fig. 5.**
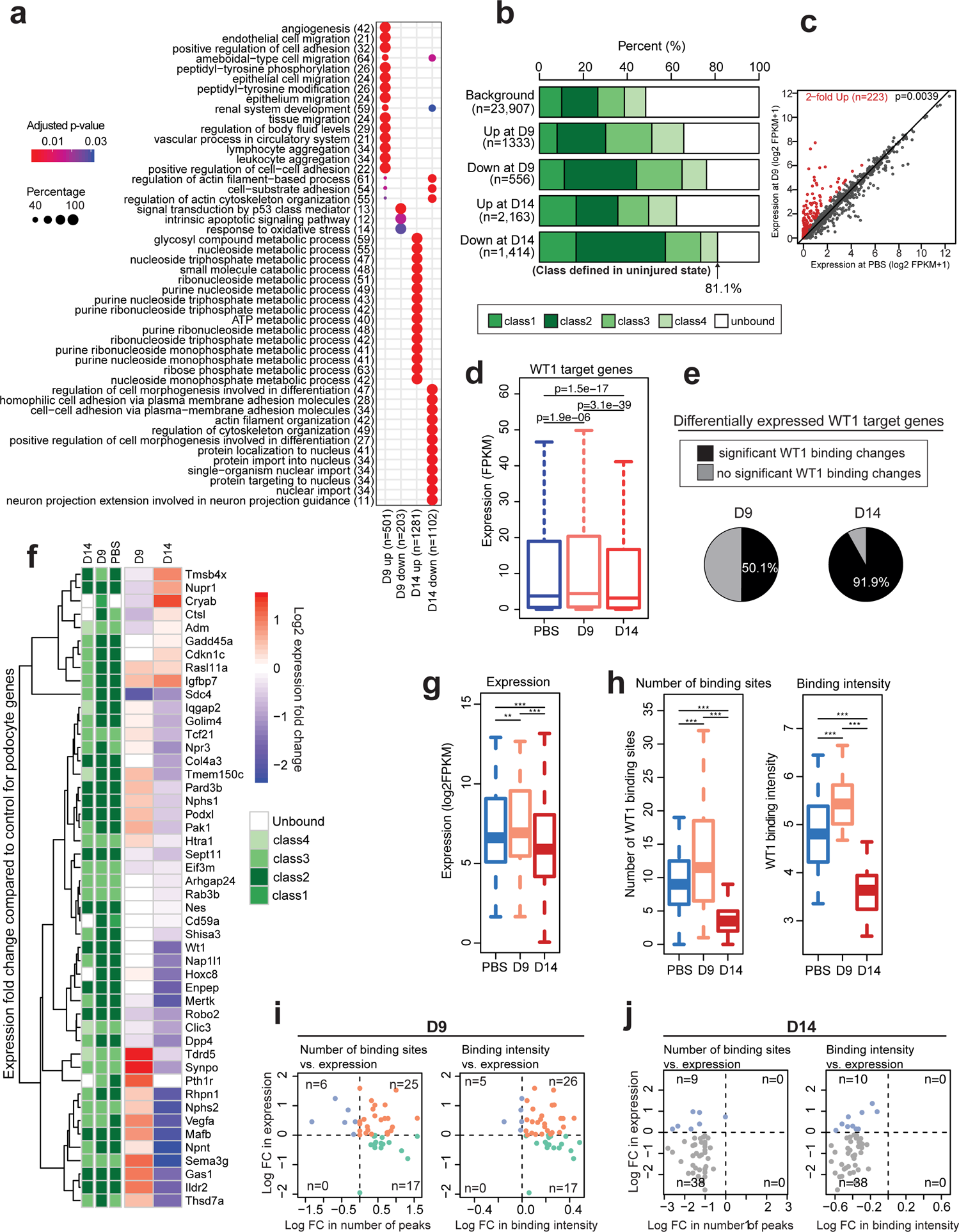
Transient increase of gene expression during ADR-induced injury in *mTmG-Nphs2cre* mice. (**a**) GO terms enriched for differentially expressed genes (see Methods). D9 Up: genes with significantly increased expression at D9 after ADR injury. D9 Down: decreased expression at D9. D14 Up: increased expression at D14. D14 Down: decreased expression at D14. Red/blue shading indicated p-value, circle size represents percent of gene set expressed. (**b**) Proportion of differentially expressed genes in each class at D9 and D14. Gene classes are based on the WT1 binding status in uninjured podocytes (as in Fig. 4). Numbers in parenthesis represent the number of genes with differential expression using relaxed thresholds (see Methods). Background: gene class distribution among all genes. White areas represent proportion of unbound genes. (**c**) Expression ranges of WT1 target genes during the course of injury. *P*-values based on one-sided paired Wilcoxon rank sum test. (**d**) Portions of WT1 target genes with significant changes in WT1 binding at D9 (50.1%) and D14 (91.9%). (**e**) Comparison of the expression levels between control podocytes and D9 for the 1,245 genes, showing significantly increased overall expression at D9 (*P*=0.0039, One-sided paired Wilcoxon test). Each dot represents a gene. Red dots: genes with more than 2-fold increased expression. (**f**) Heatmap showing expression changes and WT1 gene classes for podocyte specific genes. The red/blue range represents the expression fold change between control and D9, D14 podocytes; red-increased, blue-decreased at D9 or D14 compared to control. Green colors above heatmap indicate gene classes based on WT1 binding at each time point. (**g**) Expression range of podocytes genes during the course of injury. *P*-values by one-sided paired Wilcoxon test. ** *P<*0.01; *\*\*\* P<*0.001. (**h**) Changes in the number of WT1 binding sites (left plot) and peak intensity (right plot) during the course of injury. *P*-values based on one-sided paired Wilcoxon test. (**i**) Correlation between changes in gene expression and changes in the number of binding sites (left plot) and in the peak intensity (right plot) at D9. FC: fold change. Orange dots: genes with increased expression associated with an increase of the number of peaks or intensity. Green dots: genes with decreased expression associated with an increase of the number of peaks or intensity. Blue dots: increased expression associated with a decrease of the number of peaks or intensity. Gray dots: decreased expression associated with a decrease of the number of peaks or intensity. (**j**) Same as in (**i**) but for D14 time point.

### WT1 target gene expression changes during injury

WT1 bound genes were found both among those that showed increased and decreased expression levels, further demonstrating that WT1 is an important determinant of gene expression in podocytes. This was the case at both D9 and D14 (Fig. 5b). Class 2 and 3 genes also showed the greatest overall differences in expression levels between control and D9 or D14 (Fig. 5b). Thus, having multiple WT1 binding sites confers a greater likelihood of gene expression levels changing during the injury process.

Of the 1,245 genes uniquely bound by WT1 at D9 (Fig. 4b), 223 increased their expression by at least 2-fold (Fig. 5c and Supplementary Fig. 5c) and 68 became expressed at D9. Thus, the majority of genes uniquely bound at D9 did not show significant differential expression between control and D9 or D14, demonstrating that changes in expression are more complex than simply reflecting *de novo* or changes in WT1 binding. Examining the entire set of WT1-bound genes, there was a slight increase in transcript levels, indicating that many WT1-bound genes are not over-expressed during the response to injury (Fig. 5d). Overall, 38% and 64% of differentially expressed genes showed a change in WT1 binding intensity at D9 and D14 respectively (Supplementary Fig. 5d). However, 50% of WT1-bound genes whose expression significantly changed at D9, also showed a change in the pattern of WT1 binding. At D14, over 90% of WT1-bound genes whose expression changed showed a change in WT1 binding intensity (Fig. 5e), emphasizing the importance of WT1 binding for gene expression during injury.

### Podocyte-specific gene expression is transiently increased during ADR-induced injury

Transcriptomic data was used to analyze WT1 binding and the expression profile of a recently described podocyte-identifying gene set^11^ during injury. Most members of this set were class 2 genes. While some class 2 genes remained in the same class, several others converted to class 3 or were unbound by D14 (Fig. 5f). Expression of this gene set significantly increased at D9 (*P*<0.01) and significantly decreased at D14 (*P*<0.001; Fig. 5g), as did the average intensity and the number of WT1 peaks (Fig. 5h). Interestingly, while the majority of genes acquired additional WT1 bound sites at D9, several of these genes actually decreased their expression level (Fig. 5i), again indicating that increased number of WT1 bound sites by itself does not necessarily confer increased expression. However, by D14, we observed a stronger correlation between the changes in number of WT1 peaks and expression levels (Fig. 5j). Similar observations were made for the correlation between WT1 binding intensity and expression (Left panels: Fig. 5i,j). In contrast, within a defined set of tubule-expressed genes^11^, WT1 only bound a subset of genes. Expression of these genes generally appeared to be regulated independently of WT1 binding (Supplementary Fig. 7b), with increased expression at D14, suggesting that podocytes may be losing their identity late in the injury process.

WT1 binds at genes encoding other major TFs found in podocytes, including FOXC2, LMX1B, TCF21 and MAFB^4^; the intensity of WT1 binding at most sites greatly decreases by D14 (Supplementary Fig. 8a,b). Based on RNA-seq analysis, *Wt1, Klf6, Tcf21, Zhx2* and *Mafb* are the most highly expressed TFs in podocytes, with *Tead1, Lmx1b* and *Foxc2* expressed at lower levels. Most of these TFs expression increased at D9 and decreased at D14, indicating that the major transcriptional network in podocytes transiently increased during the injury process (Supplementary Fig. 8c). It is likely that the concerted action of several of these TFs accounts for maximal expression of *Nphs2* and *Synpo* during the process of podocyte injury.

## DISCUSSION

Previous studies have identified WT1 target genes sets in podocytes and nephron progenitor cells^3, 4, 28^. In podocytes, WT1 targets most familial nephrotic syndrome and FSGS genes^4^. (Supplementary Table S4). Additionally, among nearly 900 recently identified eQTLs for human nephrotic syndrome^29^, 318 were within 10kb of a WT1 binding site that showed either increased or decreased numbers of WT1 binding sites at D9 and D14, respectively. Among these 104 genes showed changes at both D9 and D14 (listed in Supplementary table S5 with change in class genes after ADR injury). Here, we studied how WT1 regulates gene expression during the course of Adriamycin-induced injury in both murine and human podocytes. Our study validates human kidney organoids as a model for podocyte injury. In *mTmG-Nphs2cre* mice, WT1 DNA-binding and expression of many genes important to maintaining podocyte integrity transiently increased at the onset of proteinuria, before decreasing at later stages of podocyte injury. However, among the entire set of WT1 target genes, both increased and decreased expression levels were observed upon changes in WT1 binding, suggesting that WT1 may have both activator and repressor functions. Genes whose expression changed during the course of injury were related to multiple pathways known to be important in podocytes, including extracellular-matrix genes and their integrin receptors, glomerular slit diaphragm proteins and actin-regulatory proteins. Potential binding sites for several other TFs important in podocytes were found near WT1 bound sites. Additionally, at two important target genes, *Nphs2* and *Synpo*, ADR injury resulted in the transition from open to closed chromatin state at these genes, correlating with the loss of WT1 binding and gene expression, establishing WT1 as a major regulator of gene expression in response to podocyte injury.

Podocytes have one of the most complex cell morphologies among metazoan organisms, most obvious in the elaboration of FPs that are essential for maintaining the GFB. The cytoskeletal assembly mechanisms that form, maintain and repair FPs have only recently begun to be understood^30^. Complex cytoskeletal assembly requires the conserved action of many actin binding and regulatory proteins, many of which are WT1 target genes. For a large number of these genes, WT1 binding and their expression levels were increased at D9. It is not known whether foot process morphology results from a particular combination of cytoskeletal and adhesion proteins found in podocytes, such as specific combinations of Rho GAPs and GEFs. If this is indeed found to be the case, then WT1, acting with other TFs, may provide the specificity in determining the set of cytoskeletal regulatory proteins expressed in normal podocytes and amplified in response to injury. Integrin receptors for the extracellular matrix are also integrally involved in determining cell morphology. *α*3*β*1 integrin is known to be essential for foot process formation^31^ and indeed *Itga3* and *Itgb1* are class 2 genes and among the most highly expressed integrins in podocytes (Supplementary Fig. 9). *α*V and *β*5 integrins, that form a heterodimer implicated in podocyte injury^32, 33^, are also bound by WT1 and the expression of *Igtav* increases in response to injury (Supplementary Fig. 9).

In contrast to cytoskeleton and adhesion-related genes, many genes involved in energy metabolism were found to have decreased expression levels particularly at D14. Mitochondrial damage has been shown to be a major contributor to Adriamycin induced podocyte damage^34, 35^. WT1 also binds several genes encoding components that regulate the tricarboxylic acid cycle and ATP production in mitochondria. In many cases WT1 binding was inversely correlated with gene expression, suggesting that in addition to serving as a transcriptional activator, WT1 may for distinct gene sets act in a repressor complex. Such a function has previously been demonstrated for WT1^36, 37^, however it is not known what determines an activator *versus* repressor function for WT1.

Based on our results, we suggest a model whereby WT1, along with cofactors, activates a set of TFs that may form a complex to regulate the transcription of specific podocyte genes. In injured podocytes, an early response occurs defined by an increase in WT1 DNA binding that may recruit additional enhancer elements that loop into the TSS, increase activating epigenetic marks in the vicinity of the TSS and recruit additional coactivators to increase transcription at target genes (summarized in Fig. 6). This leads to an open chromatin conformation resulting in an increase of gene expression crucial to maintain podocytes function. This attempt at repair is followed by decreased expression of WT1, that in turn results in a decrease of its binding to TF target genes. Consequently, there is decreases expression of genes required to maintain the filtration barrier resulting in foot process effacement and proteinuria. Therefore, understanding the epigenetic landscape that occurs during podocyte injury could help to identify the key epigenetic changes that lead to FSGS. These epigenetic hallmarks may serve as biomarkers of FSGS diagnosis and progression, as well as facilitate the development of novel therapeutic approaches targeting the epigenome.

**Fig. 6.**
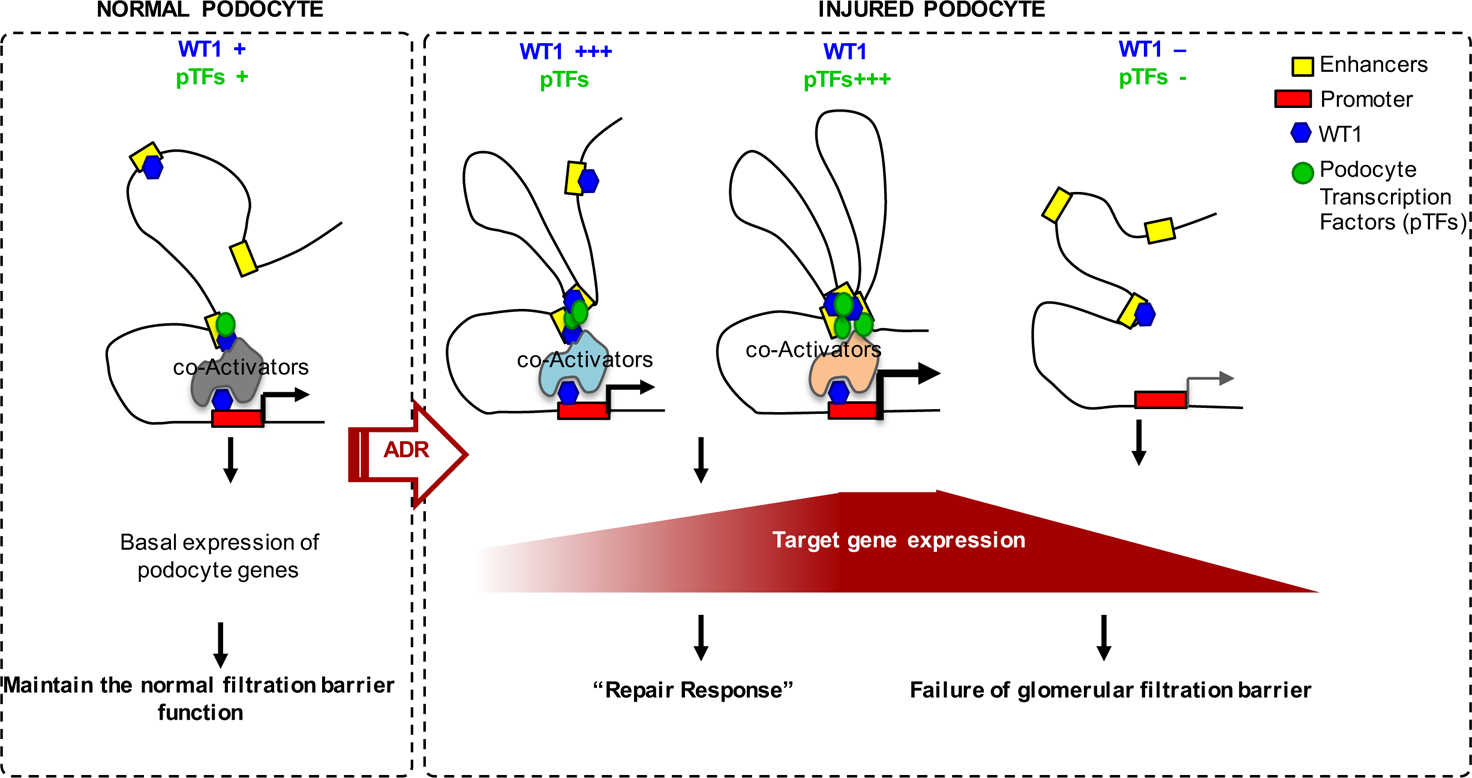
Model of WT1 transcriptional reprogramming during podocyte injury. In normal podocytes, WT1 recruits a set of TFs that form a complex to regulate the transcription of specific podocyte genes. In injured podocytes, an early response occurs by an increase in WT1 DNA binding that increases the recruitment of epigenetic coactivators. This leads to open chromatin at additional enhancers followed by increased DNA binding of additional podocyte TFs and increased transcription of podocyte-specific genes. This repair response is followed by the decreased expression of WT1 and decreased binding of WT1 and other TFs to target genes, and the inability to maintain the filtration barrier.

## METHODS

### Cell culture

Immortalized mouse podocytes were cultured with RPMI-1640 medium (Corning) with 10% fetal bovine serum, 5% sodium pyruvate solution 100 mM (Thermofisher). Undifferentiated cells were cultured at 33 °C in the presence of 10U/mL murine IFN-gamma (R&D systems). To induce podocyte differentiation cells were shifted to 37 °C for 14 days in the absence of IFN-gamma.

### Mice

All animal studies were carried out in accordance with the guidelines of the Institutional Animal Care and Use Committee at Boston Children’s Hospital. BALB/cJ mice were purchased from Jackson Laboratories. *mT/mG-Nphs2cre* mice were obtained by breeding *R26-mTmG* mice (Jackson Labs 007676) with *Nphs2cre* mice expressing red fluorescence prior to Cre recombination and green fluorescence after recombination in podocytes ^38^. This system was used to isolate podocytes by flow cytometry (FACS). BALB/cJ and *mT/mG-Nphs2cre* mice were injected with 10.5mg/kg and 18 mg/kg of Adriamycin respectively (Cayman Chemical), or PBS control, through the retro-orbital venous sinus under isoflurane anesthesia. *mT/mG-Nphs2cre* mice received two injections at a one-week interval. Kidneys were harvested at D3, D5 and D7 for BALB/cJ mice and D5, D7, D9, D10 and D14 for *mT/mG-Nphs2cre* mice. *WT1^flox/flox^/Nphs2-CreERT2/TdTomato* mice (WT1 CKO) were obtained by breeding *WT1^flox/+^* mice ^39^ with *Nphs2-CreERT2* a tamoxifen-inducible improved Cre recombinase (CreERT2) under the regulation of *Nphs2* (podocin) gene promoter ^12^ and with *R26R-tdTomato* mice (Jackson Labs 007909). This system was used to isolate tdTomato-expressing podocytes. *WT1^flox/flox^/Nphs2-CreERT2/TdTomato* mice were given tamoxifen (120mg/kg) during 3 consecutive days by intraperitoneal injections. Kidneys were harvested two weeks after the first injection.

Genotyping primers are given in supplementary material Table S1.

### Glomerular preparation and podocytes isolation

Gomerular preparation and isolation of GFP positive podocytes from 6–8 week old *mT/mG-Nphs2cre* and *WT1^flox/flox^/icre/Tdtomato* mice was done as described previously ^4^. Renal arteries were perfused with dynabeads M-450 in Hank’s balanced salt solution (HBSS), and dissected kidneys were minced and incubated in digestion solution for 15 min at 37°C (collagenase II 300 U/ml (Worthington), pronase E 1 mg/ml (Sigma), and DNAse I 50 U/μl (Applichem) in HBSS). The digest was passed through 100 μm sieves twice, washed with HBSS, spun down and glomeruli were isolated using a magnetic concentrator. Glomeruli were dissociated into a single cell suspension by incubation in digestion solution at 37°C on an incubator shaker for 40 min. Cells were sieved through a 40 μm filter and GFP-positive cells were FACS-sorted on a FACS MoFlo flow cytometer.

### Kidney organoid generation and ADR treatment

H9 human embryonic stem cells (hESCs) were differentiated into kidney organoids as previously reported ^40, 41^. Briefly, hESCs were differentiated into metanephric mesenchyme (MM) cells by a three-step directed differentiation protocol. MM cells were resuspended in 96-well, round bottom, ultra-low-attachment plates (Corning), and further differentiation was promoted by FGF9 (R&D systems) and transient treatment of CHIR (Tocris). After day 21 of differentiation, organoids were cultured in basic differentiation medium consisting of Advanced RPMI 1640 and L-GlutaMax (Life Technologies) until day 49 of differentiation. Then, kidney organoids were treated with 10μM ADR for 24hr from day 49 of differentiation. Organoids were harvested after 1, 4, 7, and 10 days of ADR injury (on day 50, 53, 56, and 59 of differentiation).

### Statistics

Two-tailed paired Student’s t-test was used to determine statistical significance between PBS and ADR conditions. Bars represent means and error bars ± SEMs. *\*\*\*P<0.001, **P<0.01; *P<0.05.* Anova with Tukey’s multiple comparisons test were used to compare different time points for WT1 ChIP-qPCR and histones ChIP-qPCR. **** *P<0.0001*, *** *P<0.001*, ** *P<0.01*, *\* P<0.05*. Multiple *t*-test with FDR determined using the two-stage linear step-up procedure of Benjamini, Krieger and Yekutieli was used to compare different conditions (PBS/ADR and control/WT1CKO) for histones ChIP-qPCR.

### Bioinformatic analysis

#### Alignments

Reads after removing adaptor sequences were mapped to mm9 using Bowtie1 ^42^ with a unique mapping option for ChIP-seq samples. Bowtie1 ^42^ and Tophat2 ^43^ were used for RNA-seq data alignment with no novel junction option.

#### Visualization of ChIP-seq tracks

Library size normalized read density profiles were generated using SPP R package get.smoothed.tag.density function ^44^ for WT1 ChIP-seq data in BALB/cJ mice. For ChIP-seq data in *mTmG-Nphs2cre* mice, aligned reads from replicates were merged after checking reproducibility for each condition. Library size normalized read density profiles were generated using deeptools bamCoverage function with RPGC normalization option and the exclusion of chrX for normalization ^45^.

#### Genomic annotations

The genomic annotations for promoters (with 500 bp margins), exons, 5’UTR, 3’UTR and genic regions were obtained from UCSC genome browser for mm9, refseq annotations. For the intergenic regions, the genic regions were subtracted from the whole genome.

#### Significant peak detection for WT1 ChIP-seq

Only uniquely aligned reads were used for downstream analysis of ChIP-seq samples. After checking reproducibility between replicates, reads in replicates were combined. Significant peaks were detected using MACS2 callpeak function with q-value=0.05 ^46^.

#### Differential bindings of WT1

For the union peaks from two conditions, read counts were obtained and statistically compared between two conditions using Diffbind R package ^47^, which internally implements Deseq2 ^48^ to access dispersion and significance of the fold changes. The WT1 changes with FDR < 0.05 and log2 fold change > 0.5 were considered to be significant changes for the downstream analysis for genomic distributions in Figures 2C and supplemental 1C and WT1 gene classes in Figures 2F, 4D and supplemental 1E.

#### Transcription factor (TF) enrichment around WT1 peaks

For the significantly increased WT1 bindings at D9 compared to control (FDR<0.05 and a fold change > 2), DNA sequences were obtained with a 200 bp window for Figure 3A. For the control sequences, the same number of WT1 binding sites as those of significantly changed ones were chosen from the locations where WT1 binding does not change significantly at D9 (p-value>0.05 and a log2 fold change < 0.5). The enrichment of TF motifs was compared between the sequences around the significant change of WT1 binding sites and those around non-changed WT1 binding sites using the MEME suite AME ^49^ with default parameters. TF motif sequences were collected from Jaspar core 2018 vertebrate database ^50^ and Jolma *et al* ^51^. Among the significantly enriched TFs, TFs that are silent or low-expressed in podocytes were filtered out. The same procedure was done for the prediction of enriched TFs at the significantly decreased WT1 binding sites at D14 in Figure 3C. For the prediction of enriched TFs for unique WT1 peaks at D9, DNA sequences were obtained with a 200 bp window from D9 unique WT1 peak summits at D9. Persistent peak sites found in PBS, D9 and D14 were used as control sites.

#### TF binding prediction near WT1 binding sites for Nphs2 and Synpo

For the WT1 binding sites overlapping with regions of high conservation scores, the locations of key TFs in podocytes such as TEAD1, FOXC2, TCF21, WT1, MAFB and LMX1B and several TFs enriched for differential WT1 binding sites during the course of ADR injury were predicted using the MEME suite FIMO with p=0.01^52^. The window size of 100 bp was used from the WT1 peak summits to extract DNA sequences. Motif database from Jaspar core 2018 vertebrate database ^50^ and Jolma *et a*l ^51^ were used.

#### Association between WT1 peaks and genes

To infer potential target genes of WT1 peaks, for the proximal regions, upstream of 5 kb and downstream of 1 kb regions of transcription start sites (TSSs) of the genes were associated with the WT1 binding site. For the distal regions, up to 500 kb regions were considered using GREAT version 2 ^53^ for most analyses. For the de novo WT1 peaks at D9, 10 kb upstream and downstream regions of TSSs were considered in Figure 4. For the analysis of expression changes for WT1 target genes in Figure 4E, a 10 kb margin was used to associate potential targets of WT1 bindings.

#### Gene classes based on WT1 binding

For each condition, gene classes were defined as in Kann *et al* ^4^ and further modified. More specifically, Class 1 gene is the gene whose promoter (+/-1 kb from TSSs) is bound by a single WT1 peak within 500 kb regions of the TSS. Class 2 gene was the gene that has multiple WT1 bindings including peaks at the promoter. Class 3 gene was the gene having multiple WT1 bindings except for a promoter. Class 4 gene was the gene having a single peak in a non-promoter region within a 500 kb window.

#### Expression quantification

To access expression levels, Cufflinks ^54^ was with default parameters from the aligned reads. The transcriptional annotations of UCSC mm9 were used. The FPKM (fragments per kilobase of transcript per million mapped reads) values for the expression levels for each gene were calculated.

#### Differentially expressed genes

Three different methods: Cuffdiff ^54^, Deseq2 ^48^ and EdgeR ^55^, were used to detect significantly expression changed genes between conditions for a stringent gene set in Figure 4B. Q-value of 0.05 was used for Cuffdiff as a threshold, p-values of 0.05 were used for Deseq2 and EdgeR. The genes detected from at least two methods were used for the gene ontology (GO) analysis. For Figure 4D, differentially expressed genes were determined by relaxed criteria, with a p-value of 0.05 from Cuffdiff.

#### GO analysis

For the significantly changed WT1 binding sites, GREAT version 3 ^53^ was used to determine GO terms associated with the WT1 binding sites with default parameters. For the genes with significant expression changes, DAVID ^56^ was used. The results were visualized using R package clusterProfiler ^57^.

## ACKNOWLEDGEMENTS

The authors thank the members of the Kreidberg and Park Laboratories for valuable comments during the course of the work, Farhard Danesh (M.D. Anderson) for Nphs2-CreERt2 mice, Vicki Huff (M.D. Anderson) for Wt1 floxed mice, Martin Kann (M.D. Cologne, Germany) and Benoit Laurent (PhD, Sherbrooke, Canada) for advice on experimental approaches. This work was supported by 1R01DK109972-01 to J.K., DP2EB029388 and U01EB028899 to R.M., R01 DK091299-01 to V.S., and a grant from the Uehara Memorial Foundation to K.H.

