## Supplemental files for "Epigenetic transcriptional reprogramming by WT1 mediates a repair response during podocyte injury"

### Supplementary Fig. 1

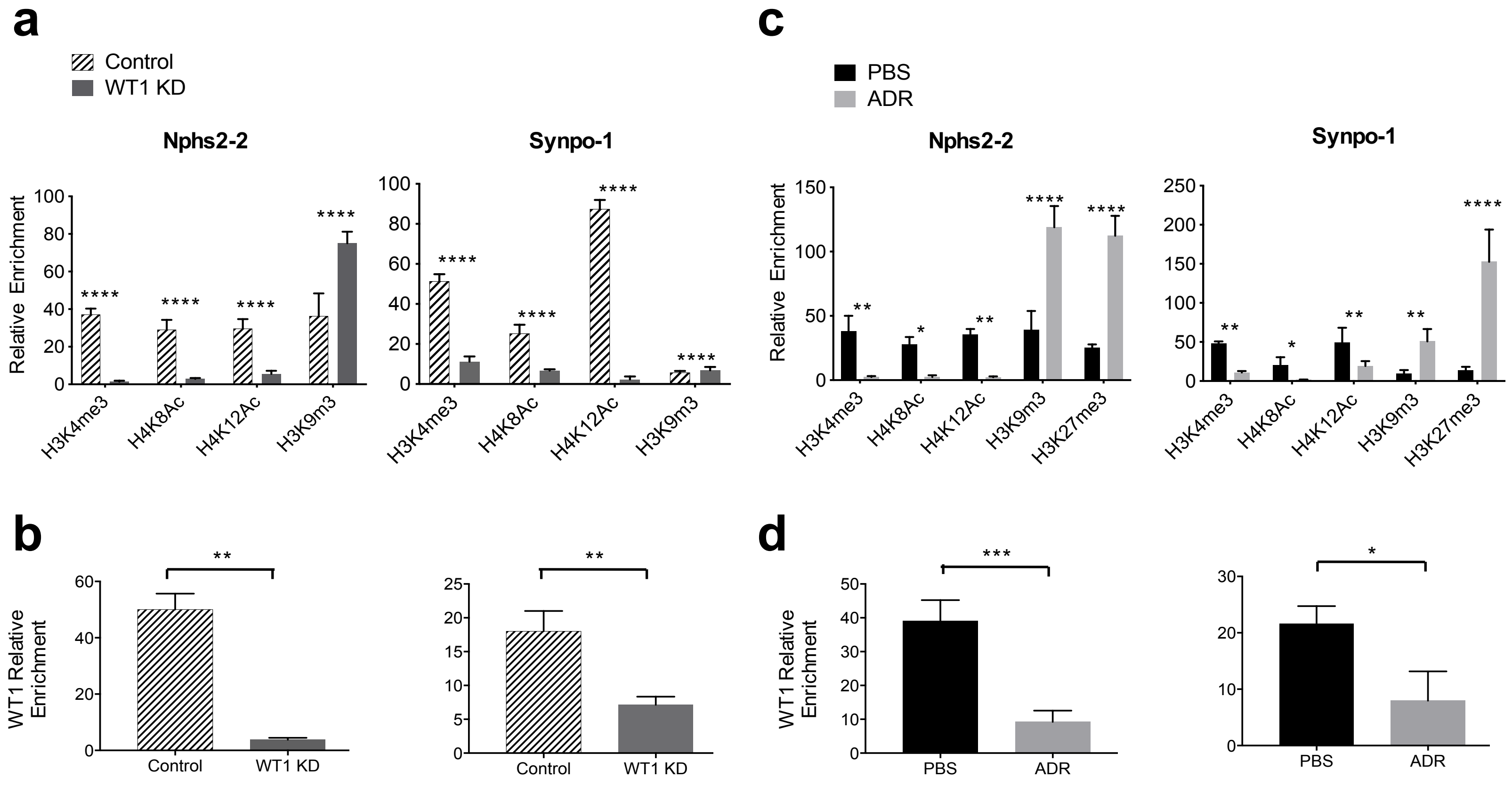

**Supplementary Fig. 1. WT1 controls chromatin remodeling at *Nphs2* and *Synpo* genes in murine immortalized podocytes**

(**a, c**) Histones direct ChIP-qPCR using active histones marks (H3K4m3, H4K8ac) and repressive histones marks (H3K9me3 and H3K27me3) at Nphs2-2 and Synpo-1 peaks from immortalized mouse podocytes treated with PBS or 1µg/mL of ADR during 16 hours (**a**) or podocytes transfected with siRNA scramble or siRNA WT1 (**c**). **** $P$<0.0001, *** $P$<0.001, ** $P$<0.01, * $P$<0.05 (Multiple $t$-test with FDR determined using the two-stage linear step-up procedure of Benjamini, Krieger and Yekutieli) compared to control. (**b, d**) WT1 direct ChIP-qPCR at Nphs2-2 and Synpo-1 peaks from immortalized mouse podocytes treated with PBS or 1µg/mL of ADR during 16 hours at Nphs2-2 and Synpo-1 peaks (**b**) or podocytes transfected with siRNA scramble or siRNA WT1 (**d**). Bars represent means and error bars ± SEMs. *** $P$<0.001; **$P$<0.01; *$P$<0.05 (n=3).

### Supplementary Fig. 2

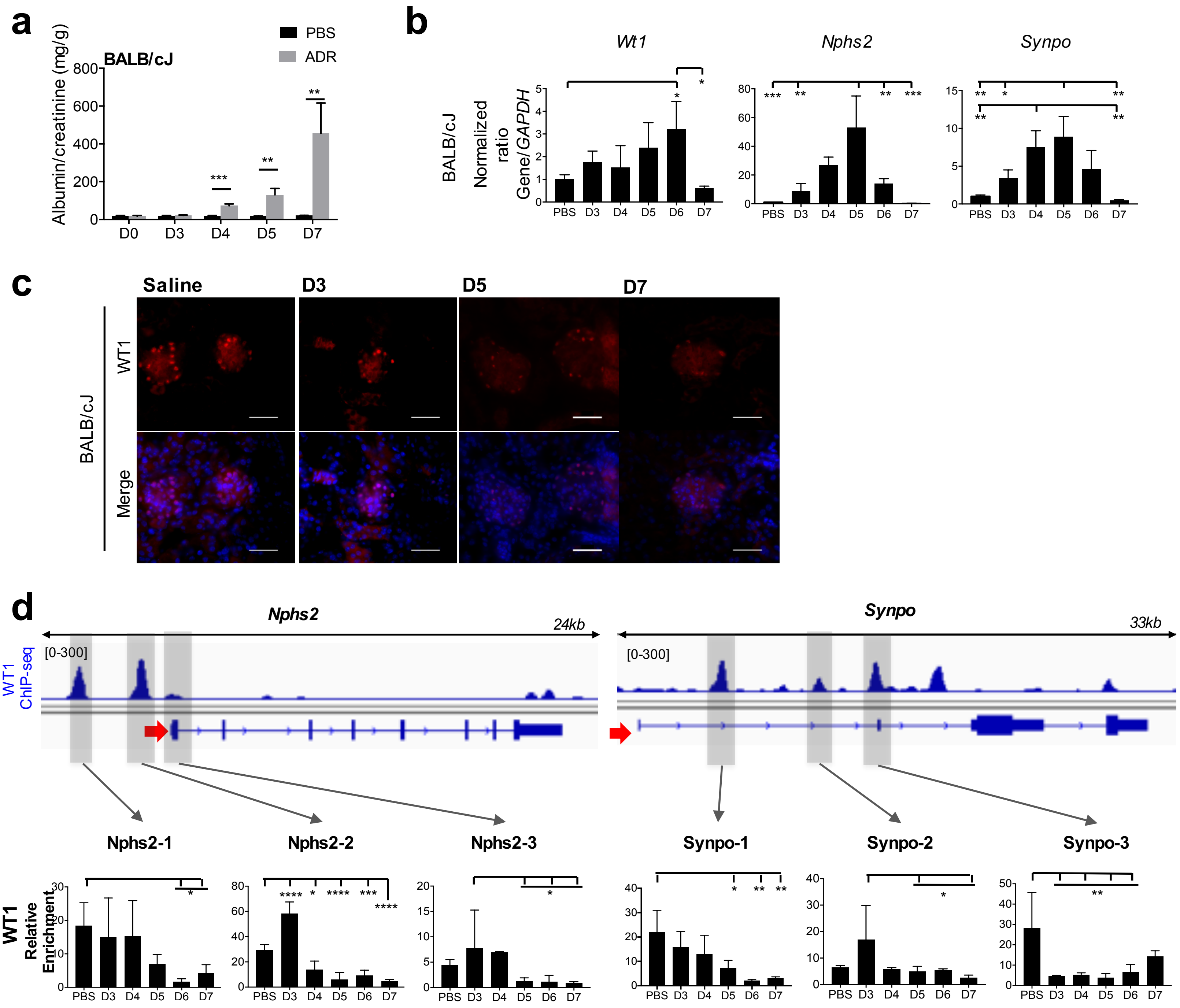

**Supplementary Fig. 2. Transient increase in the expression of key podocyte genes in ADR-injured BALB/cJ mice**

(**a**) Quantification of albumin/creatinine level during the course of ADR injury from BALB/cJ mice injected with 10.5mg/kg of ADR (grey bars) or PBS (black bars). Bars represent means and error bars ± SEMs. ****P*<0.001*;* ***P*<0.01 (n=3 replicates). (**b**) RT-qPCR of *Wt1*, *Nphs2* and *Synpo* from isolated glomeruli from BALB/cJ during injury. One-way ANOVA with Tukey's multiple comparisons test were used. ****P*<0.001; ***P*<0.01; **P*<0.05 (n=3 replicates). (**c**) Immunofluorescent staining of WT1 (red) in BALB/cJ mice in glomeruli. Scale bar 50µM. (**d**) Upper panels representing IGV plots of *Nphs2* and *Synpo* genes for WT1 ChIP-seq showing WT1 binding sites (gray highlighted boxes) in uninjured podocytes: Nphs2-1, Nphs2-2, Nphs2-3, Synpo-1, Synpo-2 and Synpo-3**.** Lower panels: WT1 dynamic binding at *Nphs2* and *Synpo* genes measured by WT1 direct ChIP-qPCR from isolated glomeruli from BALB/cJ mice (n=3). ANOVA with Tukey's multiple comparisons test were used. **** *P*<0.001; ****P*<0.001; ***P*<0.01; **P*<0.05.

### Supplementary Fig. 3

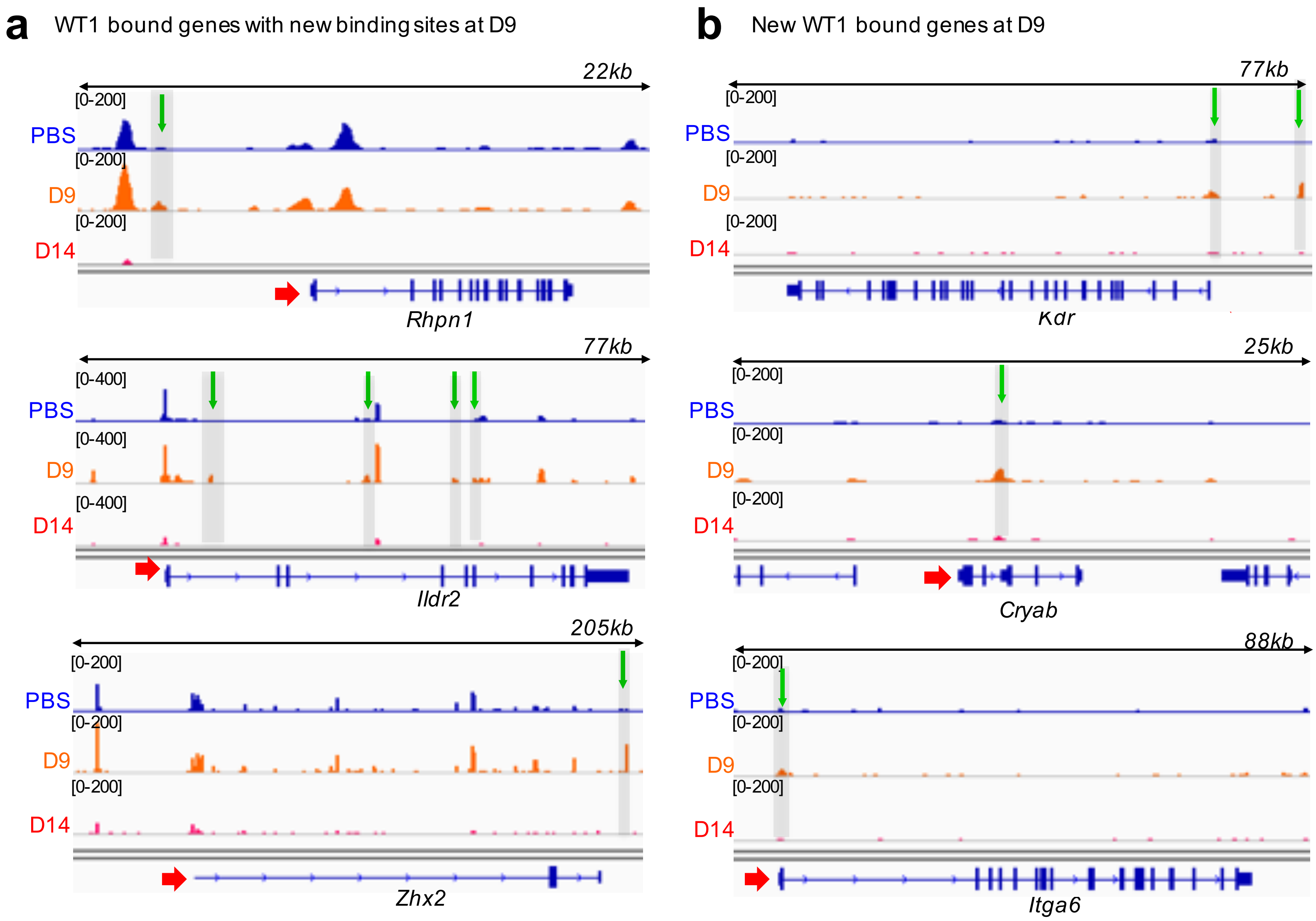

**Supplementary Fig. 3. Identification of new WT1 bound sites at the onset of proteinuria in *mTmG-Nphs2cre* mice**

(**a** and **b**) Identification of new WT1 binding sites at D9 within genes that were already bound (**a**) or unbound (**b**) in uninjured podocytes. WT1 ChIP-seq IGV plots of *Rhpn1, Ildr2 and Zhx2 genes* (**a**) or *Kdr, Cryab and Itga6* genes (**b**) showing WT1 binding sites during injury (uninjured/PBS: blue, D9: orange, D14: red). Red arrows show TSSs and transcription direction. Green arrows show new WT1 binding sites present at D9.

Supplementary fig 4

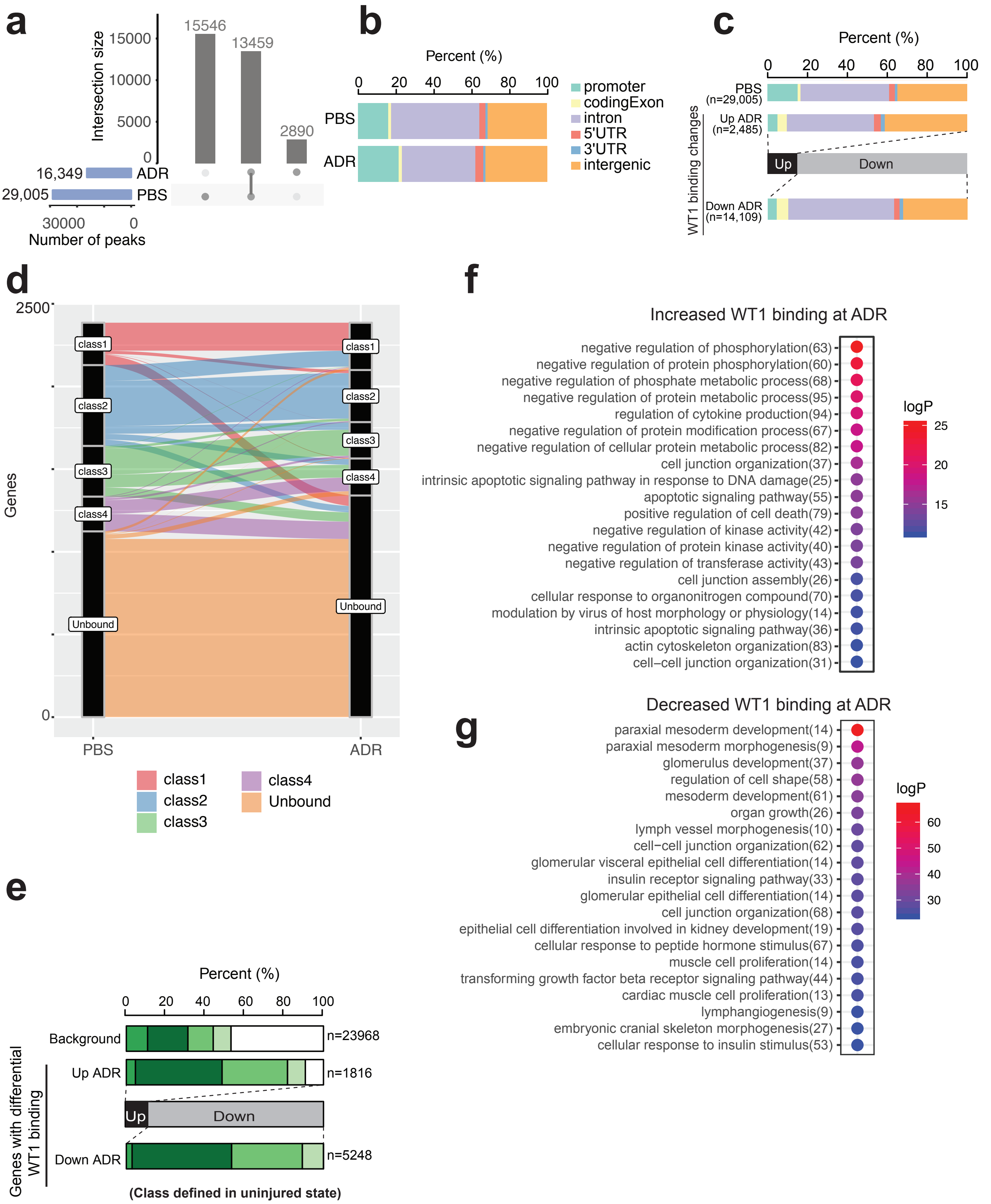

**Supplementary Fig. 4. Dynamics of WT1 binding during the injury response in BALB/cJ mice**

(**a**) Number of WT1 binding sites in control and after ADR. Grey bars represent the number of peaks common to each condition. Blue bar plot shows total binding site number. (**b** and **c**) Genomic distribution of (**b**) all WT1 binding sites or (**c**) WT1 binding sites that significantly changed during injury. (**d**) Alluvial diagram showing gene class changes after injury: class 1 (pink), class 2 (blue), class 3 (green), class 4 (purple) and unbound class (orange). Y-axis represents the number of genes per class, and X-axis the injury time points. (**e**) Proportion of each gene class for the genes associated with significant changes in WT1 binding intensity after ADR injury. Gene classes are based on the WT1 binding status in uninjured podocytes. Background indicates the distribution of gene classes for all bound genes (green). White represents unbound genes. The number on the top of the last two columns represents the number of genes with WT1 binding sites that significantly changed during the course of injury. **(f)** GO terms representing genes at which WT1 binding increased after ADR (upper panel) or decreased (lower panel).

### Supplementary Fig. 5

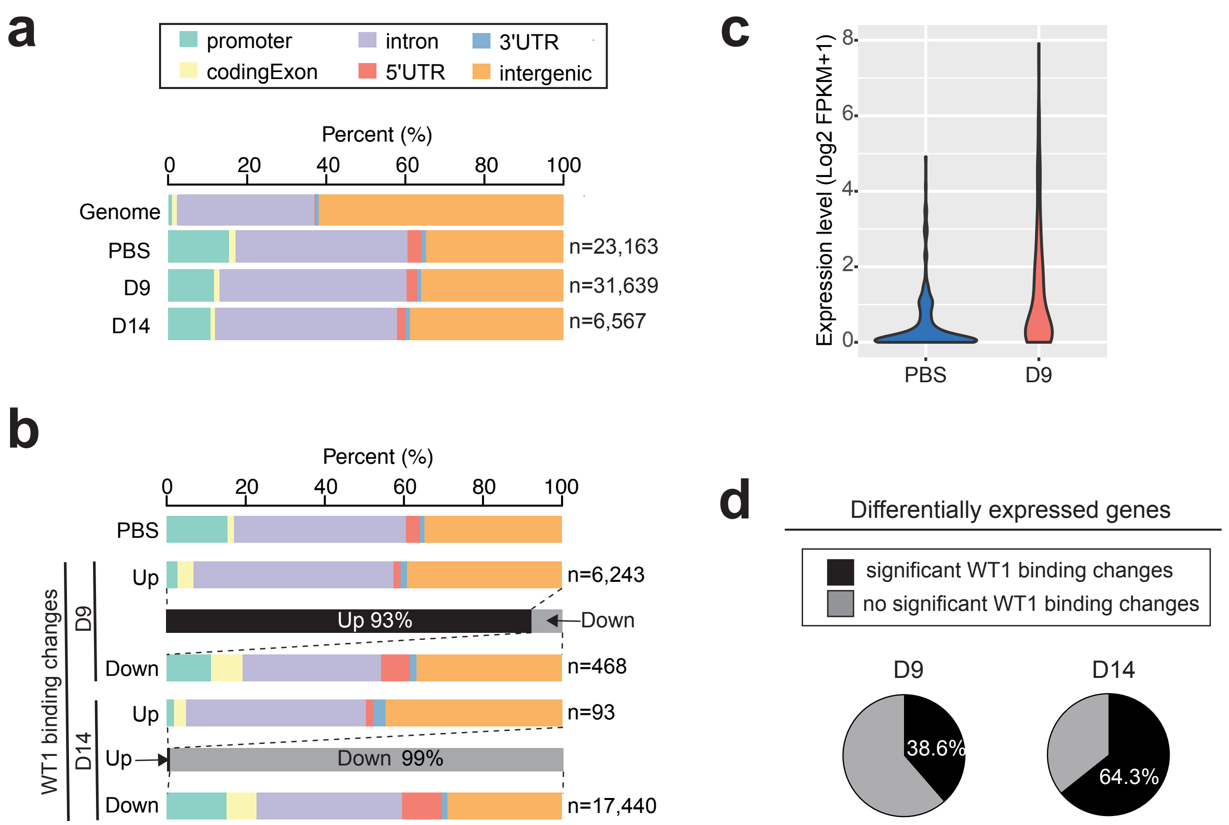

**Supplementary Fig. 5. Effect of ADR on WT1 binding and gene expression in in *mTmG-Nphs2cre* mice**

(**a** and **b**) Genomic distribution of all WT1 binding sites (**a**) or WT1 binding sites that significantly changed during injury (**b**). (**c**) Expression levels of the 223 genes with more than 2 fold increase of expression at D9 compared to control. The majority of genes were silent in uninjured podocytes. (**d**) Portions of differentially expressed genes with significant changes in WT1 binding at D9 (38.6%) and D14 (64.3%).

### Supplementary Fig. 6

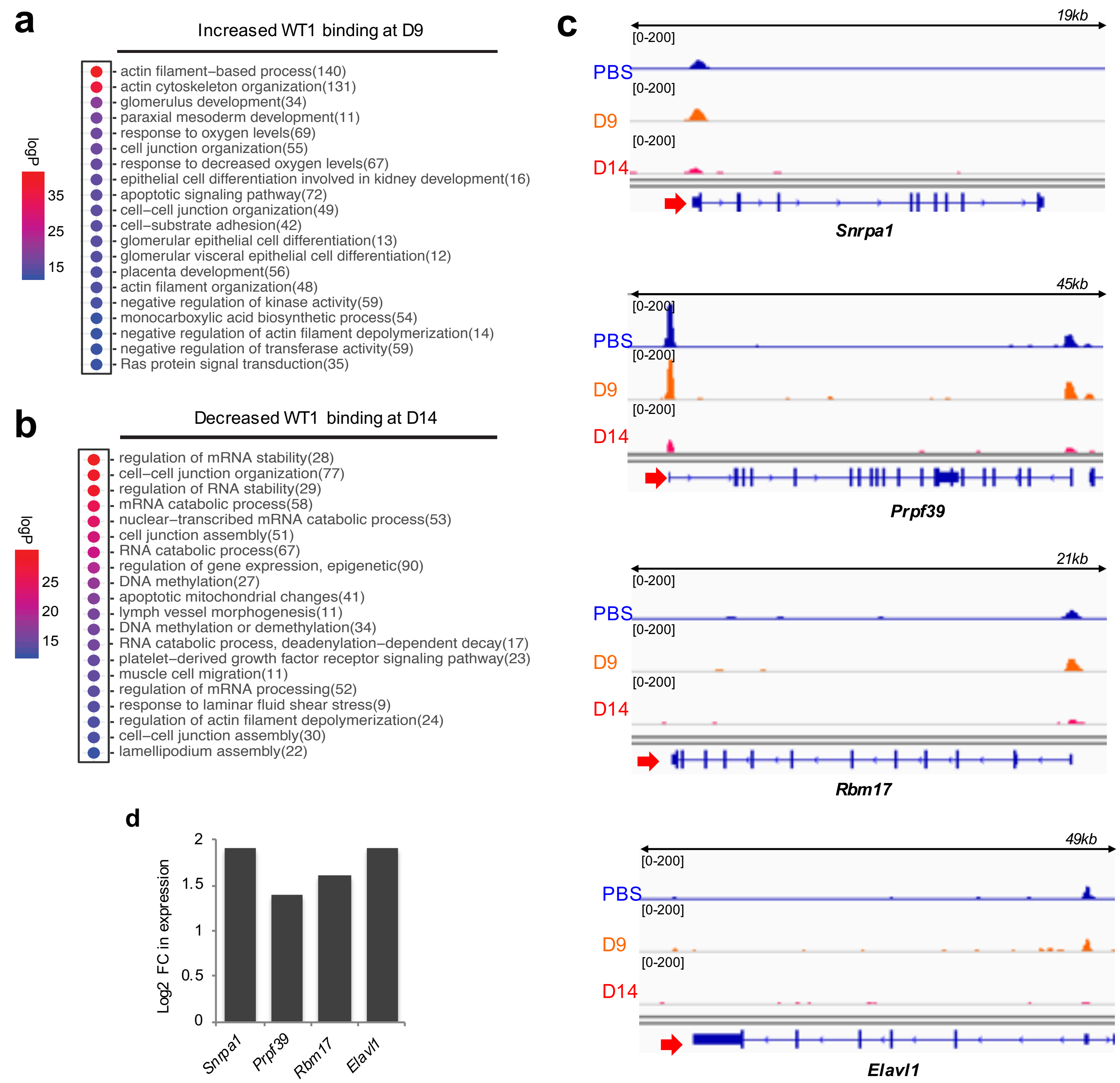

**Supplementary Fig. 6. Gene ontology terms enriched for significant WT1 binding after ADR in *mTmG-Nphs2cre* mice**

(**a**) GO terms enriched for the genes associated with significantly increased WT1 binding at D9. (**b**) GO terms enriched for the genes associated with significantly decreased WT1 binding at D14. (**c**) WT1 ChIP-seq IGV plots showing examples of increased expressed genes associated with a decrease of WT1 binding at D14. Uninjured/PBS: blue, ADR-D9: orange, ADR-D14: red. Red arrows show TSSs and transcription direction. (**d**) Log2 expression fold change of genes shown in (**c**). FC: fold change.

### Supplementary Fig. 7

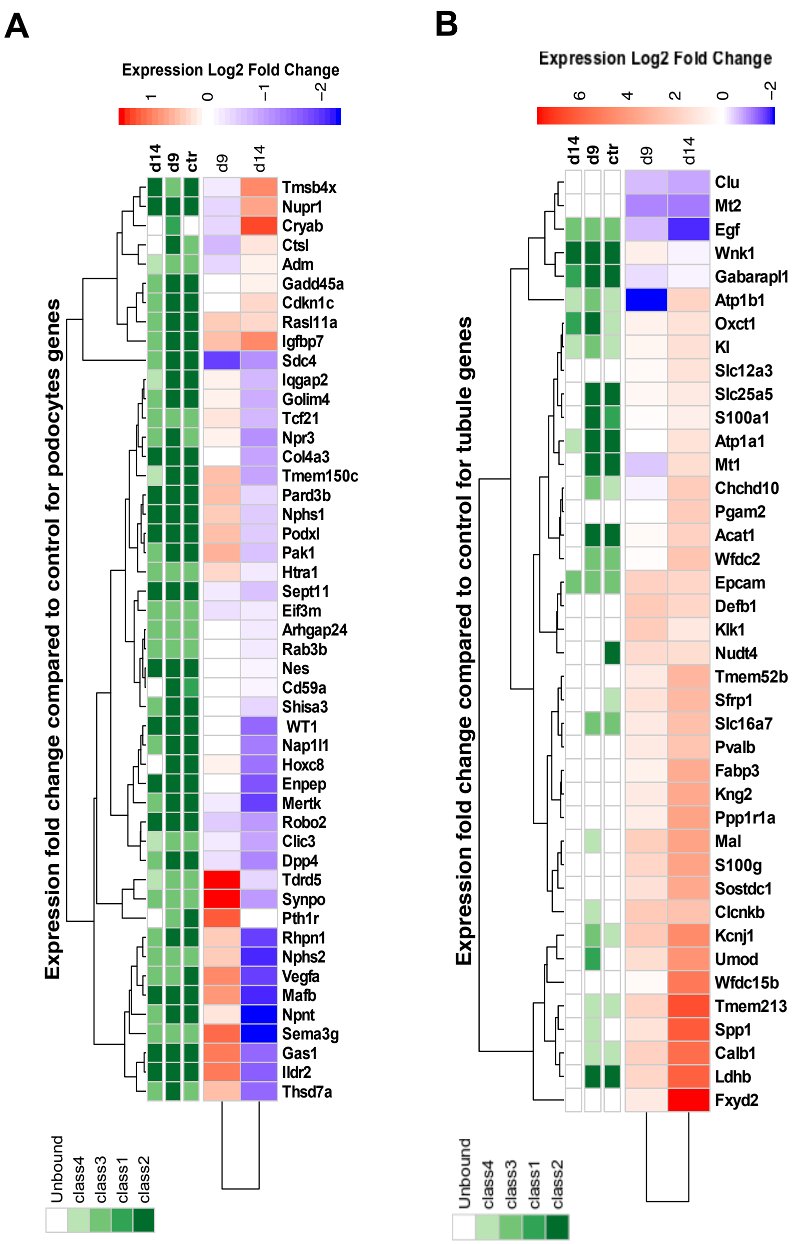

**Supplementary Fig. 7. Podocyte specific gene expression during ADR-induced injury in *mTmG-Nphs2cre* mice**

(**a** and **b**) Heatmap showing the expression changes and WT1 gene classes for podocyte specific genes (**a,** reproduced from Figure 5 for comparision) or for tubule specific genes (**b**). Red: increased expression in injured podocytes compared to uninjured podocytes. Blue: decreased expression. Colors on the top indicate gene classes based on WT1 binding status.

### Supplementary Fig. 8

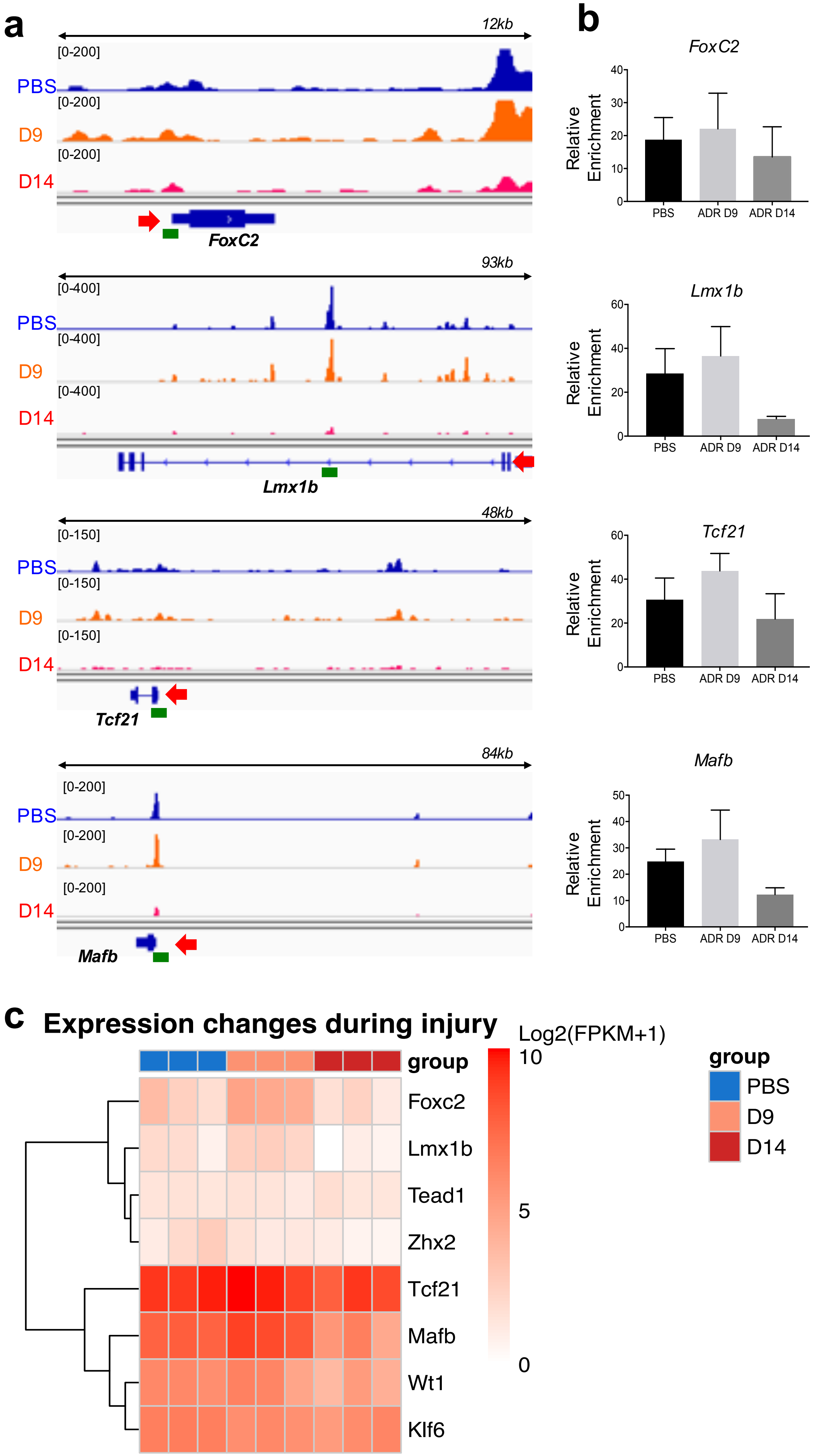

**Supplementary Fig. 8. WT1 binding and gene expression during ADR-induced injury in *mTmG-Nphs2cre* mice at podocyte TFs genes**

(**a**) WT1 ChIP-seq IGV plots of *FoxC2, Lmx1b, Tcf21 and Mafb* genes showing WT1 binding sites during injury (uninjured/PBS: blue, D9: orange, D14: red). Red arrows show TSS. Green lines show amplicon for direct ChIP-qPCR. (**b**) WT1 direct ChIP-qPCR after ADR injury from *mTmG-Nphs2cre* isolated glomeruli (n=3 replicates). (**c**) Heatmap representing the expression level of transcription factors that were found enriched at peaks with significant binding changes during injury.

**Supplementary Fig. 9**

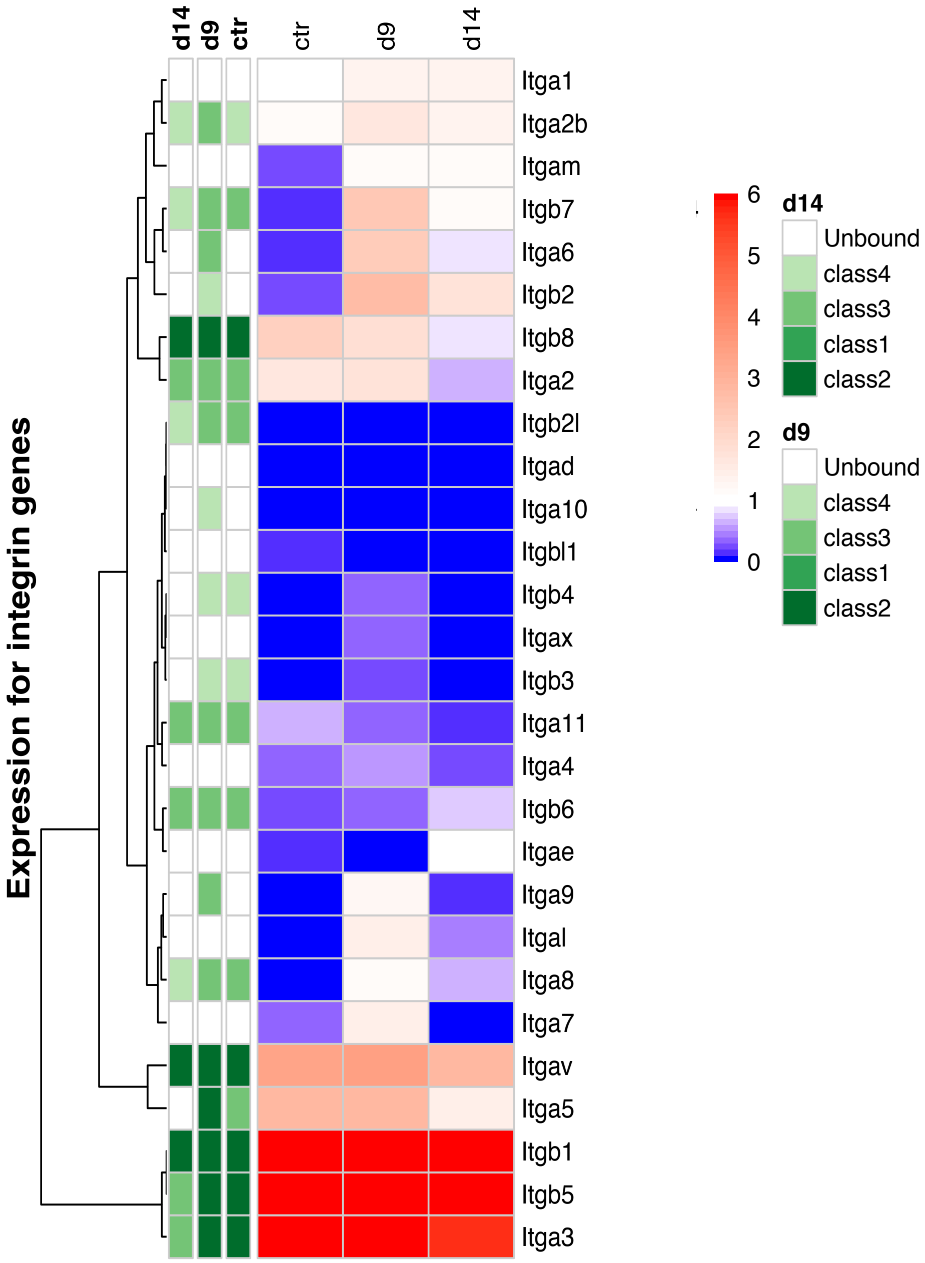

**Supplementary Fig. 9. Integrins gene expression during ADR-induced injury in *mTmG-Nphs2cre* mice**

Heatmap showing the expression changes and WT1 gene classes for integrin genes. The red/blue range represents the expression fold change between control and D9, D14 podocytes; red-increased, blue-decreased at D9 or D14 compared to control. Green colors above heatmap indicate gene classes based on WT1 binding at each time point.

#### Supplementary Table S1. Primers sequences used for genotyping

| Alleles | Sequence Primer forward (5'-3') | Sequence Primer reverse (5'-3') |
|---|---|---|
| *R26-mTmG* | CTC TGC TGC CTC CTG GCT TCT | WT-rev: CGA GGC GGA TCA CAA GCA ATA |
| | | Mutant-rev: TCA ATG GGC GGG GGT CGT T |
| *Nphs2:Cre* | GGA CAT GTT CAG GGA TCG CCA GGC G | GCA TAA CCA GTG AAA CAG CAT TGC TG |
| *WT1 flox* | CCT TTT ACT TGG ACC GTT TG | GGG GAG CCT GTT AGG GTA |
| *Nphs2:iCre* | TCA ACA TGC TGC ACA GGA GAT | ACC ATA GAT CAG GCG GTG GGT |
| *R26-TdTomato* | WT-fwd: AAG GGA GCT GCA GTG GAG TA | WT-rev: CCG AAA ATC TGT GGG AAG TC |
| | Mutant-fwd: CTG TTC CTG TAC GGC ATG G | Mutant-rev: GGC ATT AAA GCA GCG TAT CC |

#### Supplementary Table S2. Primers sequences used for RT-qPCR

| Genes | Sequence Primer forward (5'-3') | Sequence Primer reverse (5'-3') |
|---|---|---|
| *Nphs2* (m) | TGC TAC TAC CGC ATG GAA AAT G | GCA CAA CCT TTA TGC AGA ACC AG |
| *Synpo* (m) | CTG CAT CCG TGG TCA ACA G | GGG ACT CCT ATC CGC CAT AC |
| *WT1* (m) | GAG AGC CAG CCT ACC ATC C | CCC TGC TGT CCA TTC TCA AT |
| *Gapdh* (m) | GGT GAA GGT CGG TGT GAA | CAA TGA AGG GGT CGT TGA T |
| *Nphs2* (h) | AAG AGT AAT TGG ACA T | TGG TCA CGA TCT CAT GAA AAG G |
| *Synpo* (h) | CCC AAG GTG ACC CCG AAT | CTG CCG CCG CTT CTC A |
| *WT1* (h) | GTG ACT TCA AGG ACT GTG AAC G | CGG GAG AAC TTT CGC TGA CAA |
| *Nphs1* (h) | CTG CCT GAA AAC CTG ACG GT | CGA CCT GGC ACT CAT ACT CC |
| *Gapdh* (h) | GGC TCT CCA GAA CAT CAT CCC TGC | GGG TGT CGC TGT TGA AGT CAG AGG |

(m): mouse; (h): human

### Supplementary Table S3. Primers sequences used for ChIP

| Genes | Sequence Primer forward (5'-3') | Sequence Primer reverse (5'-3') |
|---|---|---|
| *Nphs2-1* | ACC TGG TCT CTT CAC AGC AC | TCC GCA GTG ACC TGG TAT TTG |
| *Nphs2-2* | TAT CCG TAA CCC CAA CCA AC | TGA GGG GGC AAA CAT TTA AG |
| *Nphs2-3* | ATC CAG ACC CAA GAA GGA AC | CCC TGC TGT CCA TTC TCA A' |
| *Synpo-1* | CCT GCC TTG AGT CCT TTC TG | CTG TTA GGG CAG AGC AGA CC |
| *Synpo-2* | TGC TGG CAC TCT GGC TAC TC | TGT GTG GGC AGC TAC TTG AG |
| *Synpo-3* | CCG ACG AAG AGA GAG GAA AA | CCG GTG AAT CTG GTG AAT CT |
| *FoxC2* | ATG TTC GAG AAT GGC AGC TT | GAC TTT CTT CTC GGC CTC CT |
| *Lmx1b* | GGC CAG AGA AGT GGG TAA CA | CTG CAA ACA CCA AGG GAA CT |
| *Tcf21* | AAA GGG TGG AGA GGG TGA GT | TGT TTC GGG GTT CCA GTT AG |
| *Mafb* | AAG GTC GAA GTC GTT GAC GTA | GTC CCC AGA CAA AGG CTT G |
| *Gapdh* | CAG GAG CCC AGG GAA GAT ACA AAT A | ACG CAT ACA CAT ATA CAA CCA GTC A |

#### Supplementary Table S4. WT1 binding status and gene expression at mutated nephropathy genes

| Gene symbol | Protein | WT1 binding | | | Expression change compared to control* | | REFs |
|---|---|---|---|---|---|---|---|
| | | Control | D9 | D14 | D9 | D14 | |
| ***ACTN4*** | *α-actinin 4* | class2 | class2 | class2 | down | down | 1 |
| ***ADAMTS9*** | *ADAM Metallopeptidase with Thrombospondin Type 1 Motif 9* | class2 | class2 | class2 | down | down | 2 |
| ***ADCK4*** | *AarF Domain Containing Kinase 4* | Unbound | Unbound | Unbound | down | down | 3 |
| ***ALG1*** | *Asparagine-linked glycosylation 1* | class2 | class2 | Unbound | up | down | 4 |
| ***ANLN*** | *Anillin actin binding protein* | Unbound | Unbound | Unbound | up | up | 5 |
| ***APOL1*** | *Apolipoprotein L1* | (no mouse homolog) | | | | | 6 |
| ***ARHGAP24*** | *Rho GTPase-activating protein 24* | class3 | class3 | class3 | no change | down | 7 |
| ***ARHGDIA*** | *Rho GDP dissociation inhibitor α* | class1 | class1 | Unbound | down | down | 8 |
| ***ARHGEF17*** | *Rho Guanine Nucleotide Exchange Factor 17* | class2 | class2 | class3 | up | down | 9 |
| ***BPTF*** | *Bromodomain PHD Finger Transcription Factor* | class2 | class2 | Unbound | up | up | 9 |
| ***CD151*** | *CD151 antigen* | class2 | class2 | class1 | down | up | 10 |
| ***CD2AP*** | *CD2-associated protein* | class2 | class2 | class1 | up | down | 11 |
| ***CDK20*** | *Cyclin-dependent kinase* | class2 | class2 | Unbound | no change | down | 12 |
| ***CFH*** | *Complement factor H* | Unbound | Unbound | Unbound | up | down | 13 |
| ***COL4A3*** | *Type IV collagen α3* | class2 | class2 | class2 | no change | down | 14 |
| ***COL4A4*** | *Type IV collagen α4* | class2 | class2 | class2 | no change | down | 14 |
| ***COQ2*** | *Coenzyme Q2* | class4 | class4 | Unbound | up | up | 15 |
| ***COQ6*** | *Coenzyme Q6* | Unbound | class4 | Unbound | down | up | 16 |
| ***CRB2*** | *Crumbs family member2* | class3 | class3 | class3 | up | no change | 17 |
| ***CUBN*** | *Cubilin* | class3 | class3 | class3 | up | up | 18 |
| ***DGKE*** | *Diacylglycerol kinase ε* | class3 | class3 | Unbound | up | down | 19,20 |

| | | | | | | | |
|---|---|---|---|---|---|---|---|
| ***DLC1*** | *DLC1 Rho GTPase-activating protein* | class3 | class3 | class3 | down | down | 12 |
| ***DLG5*** | *Discs Large MAGUK Scaffold Protein 5* | class2 | class2 | class2 | down | up | 9 |
| ***E2F3*** | *E2F transcription factor* | class2 | class2 | class3 | up | down | 21 |
| ***EMP2*** | *Epithelial membrane protein 2* | class1 | class2 | Unbound | up | up | 22 |
| ***FAT1*** | *FAT atypical cadherin 1* | class2 | class2 | class4 | up | down | 23 |
| ***GAPVD1*** | *GTPase activating protein and VPS9 domains 1* | class3 | class3 | Unbound | up | down | 24 |
| ***GCC1*** | *GRIP And Coiled-Coil Domain Containing 1* | class4 | class4 | Unbound | down | down | 9 |
| ***GPC5*** | *Glypican 5* | class3 | class3 | Unbound | down | up | 25 |
| ***INF2*** | *Inverted formin 2* | class2 | class2 | class2 | down | up | 26 |
| ***ITGA3*** | *Integrin α3* | class2 | class2 | class3 | no change | down | 27 |
| ***ITGB4*** | *Integrin β4* | class4 | class4 | Unbound | up | up | 28 |
| ***ITSN1*** | *Intersectin protein* | class2 | class2 | class2 | down | up | 12 |
| ***ITSN2*** | *Intersectin protein* | class2 | class2 | Unbound | up | no change | 12 |
| ***KANK1*** | *Kidney ankyrin repeat-containing protein* | class2 | class2 | class3 | down | down | 9 |
| ***KAT2B*** | *Lysine Acetyltransferase 2B* | class2 | class2 | class4 | up | down | 9 |
| ***LAMB2*** | *Laminin subunit β2* | class2 | class2 | class1 | no change | down | 29 |
| ***LMNA*** | *Lamin A and C* | class3 | class2 | class3 | down | down | 30 |
| ***LMX1B*** | *LIM homeobox transcription factor 1β* | class2 | class2 | class3 | up | down | 31 |
| ***MAGI2*** | *Membrane Associated Guanylate Kinase, inverted 2* | class3 | class3 | class3 | up | down | 12 |
| ***MEFV*** | *Pyrin* | Unbound | Unbound | Unbound | up | up | 32 |
| ***MYH9*** | *Myosin heavy chain 9, non-muscle* | class2 | class2 | class2 | down | down | 33,34 |
| ***MYO1E*** | *Myosin 1E* | class3 | class3 | class3 | up | down | 35 |
| ***NEIL1*** | *Nei endonuclease VIII-like 1* | Unbound | Unbound | Unbound | up | down | 36 |
| ***NPHS1*** | *Nephrin* | class2 | class2 | class2 | up | down | 37 |
| ***NPHS2*** | *Podocin* | class2 | class3 | class4 | up | down | 38 |

| | | | | | | | |
|---|---|---|---|---|---|---|---|
| ***NUP107*** | *Nuclear pore complex protein* | class3 | class3 | Unbound | no change | down | 39 |
| ***NUP133*** | *Nuclear pore complex protein* | class1 | class1 | Unbound | no change | down | 39 |
| ***NUP160*** | *Nuclear pore complex protein* | class3 | class3 | class3 | down | down | 39 |
| ***NUP85*** | *Nuclear pore complex protein* | class1 | class1 | Unbound | up | up | 39 |
| ***NXF5*** | *Nuclear RNA export Factor 5* | class4 | class4 | Unbound | down | up | 40 |
| ***OCRL1*** | *Oculocerebrorenal syndrome of Lowe* | class2 | class2 | class2 | up | down | 41 |
| ***OSGEP*** | *KEOPS complex protein* | Unbound | Unbound | Unbound | no change | up | 42 |
| ***PAX2*** | *Paired box protein 2* | class3 | class3 | Unbound | up | up | 43 |
| ***PDSS2*** | *Decaprenyl Diphosphate Synthase Subunit 2* | class2 | class2 | Unbound | no change | up | 44 |
| ***PLCE1*** | *Phospholipase C epsilon 1* | class3 | class3 | class3 | no change | down | 45,46 |
| ***PMM2*** | *Phosphomannomuta se 2* | class3 | class3 | Unbound | up | up | 47 |
| ***PODXL*** | *Podocalyxin* | class2 | class2 | class2 | up | down | 48 |
| ***PTPRO*** | *Protein-tyrosine phosphatase- RO* | class2 | class2 | class3 | up | down | 49 |
| ***SCARB2*** | *Scavenger receptor class B, member 2* | Unbound | class2 | Unbound | down | down | 50,51 |
| ***SMARCAL1*** | *SMARCA-like protein* | class4 | class2 | Unbound | up | up | 52,53 |
| ***SYNPO*** | *Synaptopodin* | class3 | class3 | class3 | up | down | 54 |
| ***TBC1D8B*** | *TBC1 domain family member 8B* | class1 | class2 | class1 | up | down | 55 |
| ***TNS2*** | *Tensin-2* | class2 | Unbound | Unbound | down | down | 12 |
| ***TP53RK*** | *KEOPS complex protein* | Unbound | Unbound | Unbound | down | down | 42 |
| ***TPRKB*** | *KEOPS complex protein* | class2 | Unbound | Unbound | down | up | 42 |
| ***TRPC6*** | *Transient receptor potential channel C6* | unbound | class2 | unbound | up | down | 56,57 |
| ***TTC21B*** | *Tetratricopeptide repeat protein 21B* | class1 | class1 | Unbound | up | down | 58 |
| ***WDR73*** | *WD repeat domain 73* | class3 | Unbound | Unbound | up | down | 59 |
| ***WNK4*** | *WNK Lysine Deficient Protein Kinase 4* | class2 | class2 | class4 | up | down | 9 |
| ***WT1*** | *Wilms Tumor 1* | class2 | class2 | class2 | up | down | 60,61 |

| | | | | | | | |
|---|---|---|---|---|---|---|---|
| ***XPO5*** | *Exportin 5* | Unbound | Unbound | Unbound | down | down | 62 |
| ***XYLT1*** | *Xylosyltransferase 1* | class2 | class2 | class1 | up | up | 9 |
| ***ZMPSTE24*** | *Zinc metallopeptidase STE24* | Unbound | class4 | Unbound | down | down | 63 |

**Supplementary Table S5. WT1 binding status at human eQTL genes with increased expression at D9 and decreased expression at D14**

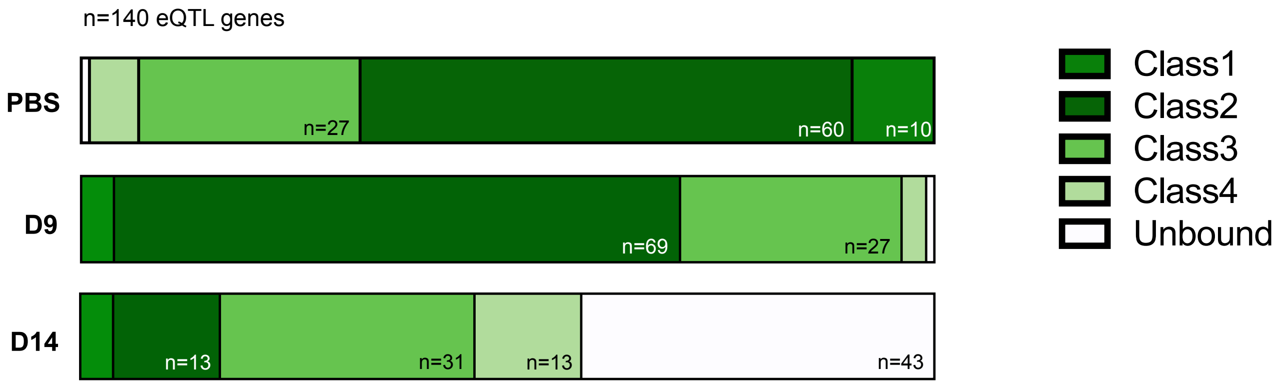


Distribution of eQTL genes by class in WT1 ChIP-seq in control (PBS), day 9 and day 14 after Adriamycin

| | | WT1 binding | | |
|---|---|---|---|---|
| **Gene symbol** | **Protein** | **Control** | **D9** | **D14** |
| ***AKAP13*** | A-Kinase Anchoring Protein 13 | class3 | class3 | class3 |
| ***ANXA5*** | Annexin A5 | class2 | class2 | class4 |
| ***APP*** | Amyloid Beta Precursor Protein | class3 | class2 | class3 |
| ***ARHGAP21*** | Rho GTPase Activating Protein 21 | class3 | class3 | class3 |
| ***ART3*** | ADP-Ribosyltransferase 3 | class2 | class3 | Unbound |
| ***ATP1B3*** | ATPase Na+/K+ Transporting Subunit Beta 3 | class2 | class2 | class4 |
| ***ATP9B*** | ATPase Phospholipid Transporting 9B | class2 | class2 | Unbound |
| ***BOK*** | BCL2 Family Apoptosis Regulator BOK | class2 | class2 | Unbound |
| ***CC2D2A*** | Coiled-Coil And C2 Domain Containing 2A | class2 | class2 | class4 |
| ***CDK5RAP2*** | CDK5 Regulatory Subunit Associated Protein 2 | class2 | class2 | class3 |
| ***CEP120*** | Centrosomal Protein 120 | class2 | class2 | class2 |
| ***CEP68*** | Centrosomal Protein 68 | class1 | class2 | Unbound |
| ***CNKSR3*** | CNKSR Family Member 3 | class2 | class2 | class4 |
| ***CNPY4*** | Canopy FGF Signaling Regulator 4 | class1 | class1 | Unbound |
| ***CREB3L2*** | CAMP Responsive Element Binding Protein 3 Like 2 | class2 | class2 | class3 |
| ***CTNNAL1*** | Catenin Alpha Like 1 | class3 | class2 | class3 |
| ***DAD1*** | Defender Against Cell Death 1 | class2 | class2 | Unbound |
| ***DCBLD2*** | Discoidin, CUB And LCCL Domain Containing 2 | class2 | class2 | class2 |
| ***DHDH*** | Dihydrodiol Dehydrogenase | class1 | class2 | Unbound |
| ***DOCK8*** | Dedicator Of Cytokinesis 8 | class2 | class3 | class3 |

| | | | | |
|---|---|---|---|---|
| ***ENOX1*** | Ecto-NOX Disulfide-Thiol Exchanger 1 | class1 | class1 | Unbound |
| ***ERAP1*** | Endoplasmic Reticulum Aminopeptidase 1 | class1 | class2 | Unbound |
| ***F2R*** | Coagulation Factor II Thrombin Receptor | class3 | class3 | class3 |
| ***FAM180A*** | Family With Sequence Similarity 180 Member A | class3 | class2 | Unbound |
| ***FAM43A*** | Family With Sequence Similarity 43 Member A | class2 | class2 | class3 |
| ***FAM81A*** | Family With Sequence Similarity 81 Member A | class3 | class3 | class3 |
| ***FDX1*** | Ferredoxin 1 | class2 | class2 | Unbound |
| ***FKTN*** | FKTN | class2 | class2 | class4 |
| ***FMO2*** | Flavin Containing Dimethylaniline Monoxygenase 2 | class1 | class3 | Unbound |
| ***GAB1*** | GRB2 Associated Binding Protein 1 | class2 | class2 | class3 |
| ***GALC*** | Galactosylceramidase | class2 | class2 | class3 |
| ***GAS7*** | Growth Arrest Specific 7 | class3 | class3 | class3 |
| ***GMEB2*** | Glucocorticoid Modulatory Element Binding Protein 2 | class1 | class2 | Unbound |
| ***HIBCH*** | 3-Hydroxyisobutyryl-CoA Hydrolase | class1 | class2 | Unbound |
| ***HTRA1*** | HtrA Serine Peptidase 1 | class3 | class3 | class3 |
| ***IFITM2*** | Interferon Induced Transmembrane Protein 2 | Unbound | class1 | Unbound |
| ***IL1R1*** | Interleukin 1 Receptor Type 1 | class2 | class2 | Unbound |
| ***IQGAP1*** | IQ Motif Containing GTPase Activating Protein 1 | class2 | class2 | class2 |
| ***IQSEC2*** | IQ Motif And Sec7 Domain ArfGEF 2 | class4 | class3 | Unbound |
| ***ITGAV*** | Integrin Subunit Alpha V | class2 | class2 | class2 |
| ***ITGB5*** | Integrin Subunit Beta 5 | class2 | class2 | class3 |
| ***ITPR1*** | Inositol 1,4,5-Trisphosphate Receptor Type 1 | class2 | class2 | class2 |
| ***KANSL1*** | KAT8 Regulatory NSL Complex Subunit 1 | class2 | class2 | class1 |
| ***KCDN3*** | Potassium Voltage-Gated Channel Subfamily D Member 3 | class2 | class2 | Unbound |
| ***KDM2B*** | Lysine Demethylase 2B | class2 | class2 | class4 |
| ***KIF16B*** | Kinesin Family Member 16B | class2 | class2 | Unbound |
| ***KIF5C*** | Kinesin Family Member 5C | class3 | class3 | Unbound |
| ***LYN1*** | LYN Proto-Oncogene, Src Family Tyrosine Kinase | class2 | class2 | class4 |
| ***LZTS2*** | Leucine Zipper Tumor Suppressor 2 | class2 | class2 | Unbound |
| ***MAP3K5*** | Mitogen-Activated Protein Kinase Kinase Kinase 5 | class2 | class2 | class3 |
| ***MCC*** | MCC Regulator Of WNT Signaling Pathway | class3 | class3 | class3 |
| ***MGAT4A*** | Alpha-1,3-Mannosyl-Glycoprotein 4-Beta-N-Acetylglucosaminyltransferase A | class3 | class3 | class4 |
| ***MICAL3*** | Microtubule Associated Monooxygenase, Calponin And LIM Domain Containing 3 | class3 | class3 | class3 |
| ***MPI*** | Mannose Phosphate Isomerase | class3 | class3 | Unbound |
| ***MRVI1*** | Murine Retrovirus Integration Site 1 Homolog | class3 | class3 | class3 |
| ***MYH9*** | Myosin Heavy Chain 9 | class2 | class2 | class2 |

| | | | | |
|---|---|---|---|---|
| ***MYOF*** | Myoferlin | class2 | class2 | class2 |
| ***N6AMT1*** | N-6 Adenine-Specific DNA Methyltransferase 1 | class2 | class2 | Unbound |
| ***NMNAT3*** | Nicotinamide Nucleotide Adenylyltransferase 3 | class1 | class4 | Unbound |
| ***PAK1*** | P21 (RAC1) Activated Kinase 1 | class2 | class2 | class3 |
| ***PAPSS2*** | 3'-Phosphoadenosine 5'-Phosphosulfate Synthase 2 | class4 | class4 | Unbound |
| ***PCSK6*** | Proprotein Convertase Subtilisin/Kexin Type 6 | class2 | class2 | class3 |
| ***PDCD1LG2*** | Programmed Cell Death 1 Ligand 2 | class3 | class3 | Unbound |
| ***PHTF1*** | Putative Homeodomain Transcription Factor 1 | class2 | class3 | Unbound |
| ***PIH1D1*** | PIH1 Domain Containing 1 | class4 | class2 | Unbound |
| ***PIP5K1B*** | Phosphatidylinositol-4-Phosphate 5-Kinase Type 1 Beta | class4 | class2 | Unbound |
| ***PKN2*** | Protein Kinase N2 | class2 | class2 | Unbound |
| ***PLA2R1*** | Phospholipase A2 Receptor 1 | class3 | class3 | Unbound |
| ***PNRC2*** | Proline Rich Nuclear Receptor Coactivator 2 | class2 | class2 | class2 |
| ***PPAT*** | Phosphoribosyl Pyrophosphate Amidotransferase | class1 | class1 | class1 |
| ***PRKCI*** | Protein Kinase C Iota | class2 | class2 | class2 |
| ***PRKG2*** | Protein Kinase CGMP-Dependent 2 | class2 | class2 | class3 |
| ***PTH1R*** | Parathyroid Hormone 1 Receptor | class2 | class3 | Unbound |
| ***RAB31*** | RAB31, Member RAS Oncogene Family | class2 | class2 | Unbound |
| ***RABGAP1L*** | RAB GTPase Activating Protein 1 Like | class2 | class2 | Unbound |
| ***RIN2*** | Ras And Rab Interactor 2 | class3 | class3 | class3 |
| ***RNF111*** | Ring Finger Protein 111 | class2 | class2 | Unbound |
| ***RNF150*** | Ring Finger Protein 150 | class2 | class2 | class3 |
| ***RSU1*** | Ras Suppressor Protein 1 | class3 | class2 | class3 |
| ***SASH1*** | SAM And SH3 Domain Containing 1 | class2 | class2 | class3 |
| ***SCML4*** | Scm Polycomb Group Protein Like 4 | class3 | class3 | Unbound |
| ***SDC4*** | Syndecan 4 | class2 | class2 | class3 |
| ***SFT2D1*** | SFT2 Domain Containing 1 | class2 | class2 | Unbound |
| ***SIDT2*** | SID1 Transmembrane Family Member 2 | class2 | class2 | Unbound |
| ***SKAP1*** | Src Kinase Associated Phosphoprotein 1 | class4 | class3 | Unbound |
| ***SLC24A3*** | Solute Carrier Family 24 Member 3 | class2 | class2 | Unbound |
| ***SLC25A39*** | Solute Carrier Family 25 Member 39 | class2 | class2 | class1 |
| ***SLC44A2*** | Solute Carrier Family 44 Member 2 | class4 | class4 | Unbound |
| ***SLC5A26*** | Solute Carrier Family 5 Member 26 | class3 | class3 | Unbound |
| ***SLCA12*** | Solute Carrier Family | class2 | class2 | class2 |
| ***SORCS2*** | Sortilin Related VPS10 Domain Containing Receptor 2 | class3 | class3 | class4 |
| ***SPINT2*** | Serine Peptidase Inhibitor, Kunitz Type 2 | class2 | class2 | class4 |
| ***ST6GALNAC3*** | ST6 N-Acetylgalactosaminide Alpha-2,6-Sialyltransferase 3 | class3 | class3 | class3 |
| ***SUCLG2*** | Succinate-CoA Ligase GDP-Forming Subunit Beta | class2 | class2 | class3 |

| | | | | |
|---|---|---|---|---|
| ***SULF2*** | Sulfatase 2 | class2 | class2 | class2 |
| ***TDRD5*** | Tudor Domain Containing 5 | class3 | Unbound | Unbound |
| ***THSD7A*** | Thrombospondin Type 1 Domain Containing 7A | class3 | class2 | class3 |
| ***TMEM135*** | Transmembrane Protein 135 | class3 | class2 | class4 |
| ***TMEM178*** | Transmembrane Protein 178A | class2 | class2 | class4 |
| ***TNIK*** | TRAF2 And NCK Interacting Kinase | class3 | class2 | class4 |
| ***UCP2*** | Uncoupling Protein 2 | class2 | class2 | class1 |
| ***VAMP8*** | Vesicle Associated Membrane Protein 8 | class2 | class2 | class2 |
| ***VEGFA*** | Vascular Endothelial Growth Factor A | class2 | class3 | class3 |
| ***WIPF3*** | WAS/WASL Interacting Protein Family Member 3 | class2 | class2 | class2 |
